# Rapid and iterative genome editing in the zoonotic malaria parasite *Plasmodium knowlesi*: New tools for *P. vivax* research

**DOI:** 10.1101/590976

**Authors:** Franziska Mohring, Melissa N. Hart, Thomas A. Rawlinson, Ryan Henrici, James A. Charleston, Ernest Diez Benavente, Avnish Patel, Joanna Hall, Neil Almond, Susana Campino, Taane G. Clark, Colin J. Sutherland, David A. Baker, Simon J. Draper, Robert W. Moon

**Author notes:** Corresponding author: Robert W. Moon, London School of Hygiene & Tropical Medicine, London WC1E 7HT.

## Abstract

Tackling relapsing *Plasmodium vivax* and zoonotic *Plasmodium knowlesi* infections is critical to reducing malaria incidence and mortality worldwide. Understanding the biology of these important and related parasites was previously constrained by the lack of robust molecular and genetic approaches. Here, we establish CRISPR-Cas9 genome editing in a culture-adapted *P. knowlesi* strain and define parameters for optimal homology-driven repair. We establish a scalable protocol for the production of repair templates by PCR and demonstrate the flexibility of the system by tagging proteins with distinct cellular localisations. Using iterative rounds of genome-editing we generate a transgenic line expressing *P. vivax* Duffy binding protein (PvDBP), a lead vaccine candidate. We demonstrate that PvDBP plays no role in reticulocyte restriction but can alter the macaque/human host cell tropism of *P. knowlesi*. Critically, antibodies raised against the *P. vivax* antigen potently inhibit proliferation of this strain, providing an invaluable tool to support vaccine development.

## Introduction

Malaria remains a serious health burden globally, with over 216 million cases annually (1). *Plasmodium falciparum* is responsible for 99 % of estimated malaria cases in sub-Saharan Africa. Outside Africa, *P. vivax* is the predominant parasite and causes ~7.4 million clinical cases annually (1). Despite extensive efforts, in 2016 the number of malaria cases were on the rise again for the first time in several years. Achieving global malaria eradication requires new tools and approaches for addressing emerging drug resistance, relapsing *P. vivax* infections, and emerging zoonotic *P. knowlesi* infections, which represent significant causes of severe disease and death (2).

Although *P. vivax* displays some distinctive features to *P. knowlesi*, including the formation of latent hypnozoites stages in the liver and restriction to reticulocytes in the blood (3), the two parasites are closely related, occupying a separate simian parasite clade to *P. falciparum* (4) Host cell invasion by *P. vivax* and *P. knowlesi* relies on the Duffy binding proteins (DBP) PvDBP and PkDBPα, respectively, both ligands for human red blood cell (RBC) Duffy antigen/receptor for chemokines (DARC) (5–8). The critical binding motif of the ligands is the cysteine-rich region 2 (DBP-RII), with ~70 % identity between PkDBPα and PvDBP (9). Despite their similarity, PvDBP has also been implicated in both *P. vivax* reticulocyte restriction (10) and as a host tropism factor preventing *P. vivax* from infecting macaques (11). PvDBP-RII is also the leading blood stage vaccine candidate for *P. vivax* (12–14), with antibodies targeting PvDBP-RII blocking parasite invasion in ex vivo *P. vivax* assays (15). *P. knowlesi* additionally contains two PkDBPα paralogues, namely DBPβ and DBPγ which share high levels of amino acid identity (68-88 %) to PkDBPα but bind to distinct receptors via N-glycolylneuraminic acid - a sialic acid found on the surface of macaque RBCs, but absent from human RBCs (16).

Due to the lack of a long-term in vitro culture system for *P. vivax*, vaccine development currently relies on recombinant protein assays, or low throughput ex vivo studies, primate infections or controlled human malaria infections (15, 17-19). Thus, higher throughput parasitological assays to assess antisera and antigens, prior to escalation to in vivo work, are desperately needed. The evolutionary similarity between *P. vivax* and *P. knowlesi* means the adaptation of *P. knowlesi* to long-term culture in human RBCs (20, 21) provides unique opportunities to study DARC-dependent invasion processes in both species. While adaptation of the CRISPR-Cas9 genome editing system to the most prevalent malaria parasite, *P. falciparum* (22), provided a powerful tool of studying parasite biology, scalable approaches for *P. falciparum* remain constrained by inefficient transfection and very high genome AT-content (averaging 80.6 %) (23). *P. knowlesi* offers significant experimental advantage*s* over *P. falciparum* including a more balanced genome AT-content of 62.5 % and orders-of-magnitude-more-efficient transgenesis (20, 24, 25).

Here, we establish CRISPR-Cas9 genome editing in *P. knowlesi*. Using an optimised and scalable PCR-based approach for generating targeting constructs we define critical parameters determining effective genome editing and apply the technique to introduce epitope/fluorescent protein tags to a variety of proteins with distinct cellular locations. We then use these tools to replace the *P. knowlesi* PkDBPα gene with its PvDBP orthologue, and delete the *P. knowlesi* DBP paralogues to create a transgenic *P. knowlesi* line reliant on the PvDBP protein for invasion of RBCs. The additional deletion of the PkDBP paralogues not only excludes interference through antibody cross-reactivity during growth inhibition assays, but also allows us to demonstrate that, in contrast to previous findings (10), PvDBP plays no role in reticulocyte restriction and has an effect on macaque/human host cell tropism. Finally, we show that antibodies raised against the *P. vivax* antigen are potent inhibitors of *P. knowlesi/* PvDBP transgenic parasites, providing an invaluable tool to support *P. vivax* vaccine development. Thus, we have developed a robust and flexible system for genome editing in an important human malaria parasite and generated essential new tools to accelerate both basic and applied malaria research.

## Results

### Homology mediated CRISPR-Cas9 genome editing is highly efficient in *P. knowlesi*

*Plasmodium* parasites lack a canonical non-homologous end joining pathway, instead relying almost exclusively on homology-directed repair of double-stranded breaks (DSBs) (26), such as those introduced by the Cas9 endonuclease. Effective CRISPR-Cas9 genome editing of malaria parasites therefore requires expression cassettes for the guide RNA and the Cas9 nuclease, and a DSB repair template (donor DNA) containing the desired change, flanked by two regions of homology to the genomic target. Whilst a variety of approaches have been used in *P. falciparum*, many embed these elements into two plasmids, each expressing a different drug-selectable marker (22, 27-30). This allows for selection of very rare events, but complicates construct design and is not ideal for multiple modifications of a given line – as both selectable markers must then be recycled. As transfection efficiency is significantly higher in *P. knowlesi* than *P. falciparum* (24), we reasoned that we may be able to use a single positive drug selectable marker to cover all the required components for editing. Pairing the guide and Cas9 cassette on a single “suicide” plasmid (29) with positive and negative selection cassettes would allow for indirect selection of a separate plasmid containing the repair template, as only parasites that took up the repair template as well as the Cas9 plasmid would be able to repair the DSB. Supporting this approach, co-transfection of plasmids expressing eGFP or mCherry revealed that ~30 % of *P. knowlesi* parasites took up both plasmids, although the proportion expressing both declined rapidly in the following days (S1 Figure). Our two-plasmid CRISPR-Cas9 system comprises one plasmid (pCas/sg) that provides Cas9, sgRNA (driven by the PkU6 promoter) and a hDHFR-yFCU fusion product for positive/negative selection, and a second plasmid (pDonor) providing the donor DNA with homology regions (HRs) flanking the DSB for repair by homologous recombination. To test this system, we designed constructs to integrate an eGFP expression cassette into the non-essential *p230p* locus (Figure 1A). A 20 bp guide sequence targeting a seed sequence upstream of a protospacer adjacent motif (PAM) within the *p230p* was cloned into pCas/sg (pCas/sg_*p230p*), and a repair template plasmid was synthesized by including an eGFP expression cassette flanked by 400 bp HRs targeting either side of the PAM sequence (pDonor_*p230p*). Both plasmids (each 20 μg) were combined and introduced into *P. knowlesi* schizonts via electroporation (20) along with control transfections (pCas9/sg without guide sequence and repair template). To simplify synchronisation of parasites, the transfection procedure was altered to additionally include a 2-hour incubation of purified schizonts with 1 μM of the schizont egress inhibitor compound 2, immediately prior to transfection. This compound reversibly inhibits the cGMP-dependent protein kinase (PKG) (31) and facilitates accumulation of the fully segmented forms required for transfection. Parasites were placed under selection with pyrimethamine for 5 days after transfection and successful integration monitored by PCR. Correct integration at the *p230p* locus was detectable by PCR within 3 days of transfection and only low levels of wild type DNA was detectable after day 11 (Figure 1B). Expression of eGFP was confirmed by live microscopy (Figure 1C). The eGFP positivity rate was calculated the day after transfection (day 1), to evaluate transfection efficiency (8.4 % ± 2.1 SD). The eGFP positivity was then assessed again once parasites reached 0.5 % parasitemia (day 12), indicating 83.3 % (± 1.8 SD) of the parasites had integrated the construct (Figure 1D). Parasites transfected with pCas/sg_*p230p* without providing pDonor_*p230p* were visible in culture several days after the integrated lines. An intact guide and PAM site was detected in these parasites, suggesting that a small population of parasites did not form DSB. Parasites transfected with pCas/sg without a cloned sgRNA appeared in culture within a few days after transfection, with comparable growth rates to the eGFP plasmid, suggesting the Cas9 expression without a targeting sgRNA is not toxic (Figure 1E). Integrated lines were grown for one week before negative selection with 5-Fluorocytosine and subsequent limiting dilution cloning. Clones were identified using a plaque-based assay (S1B Figure) previously used for *P. falciparum* (32), and 10/10 genotyped clones harboured correctly integrated, markerless eGFP (Figure 1F).

**Figure 1:**
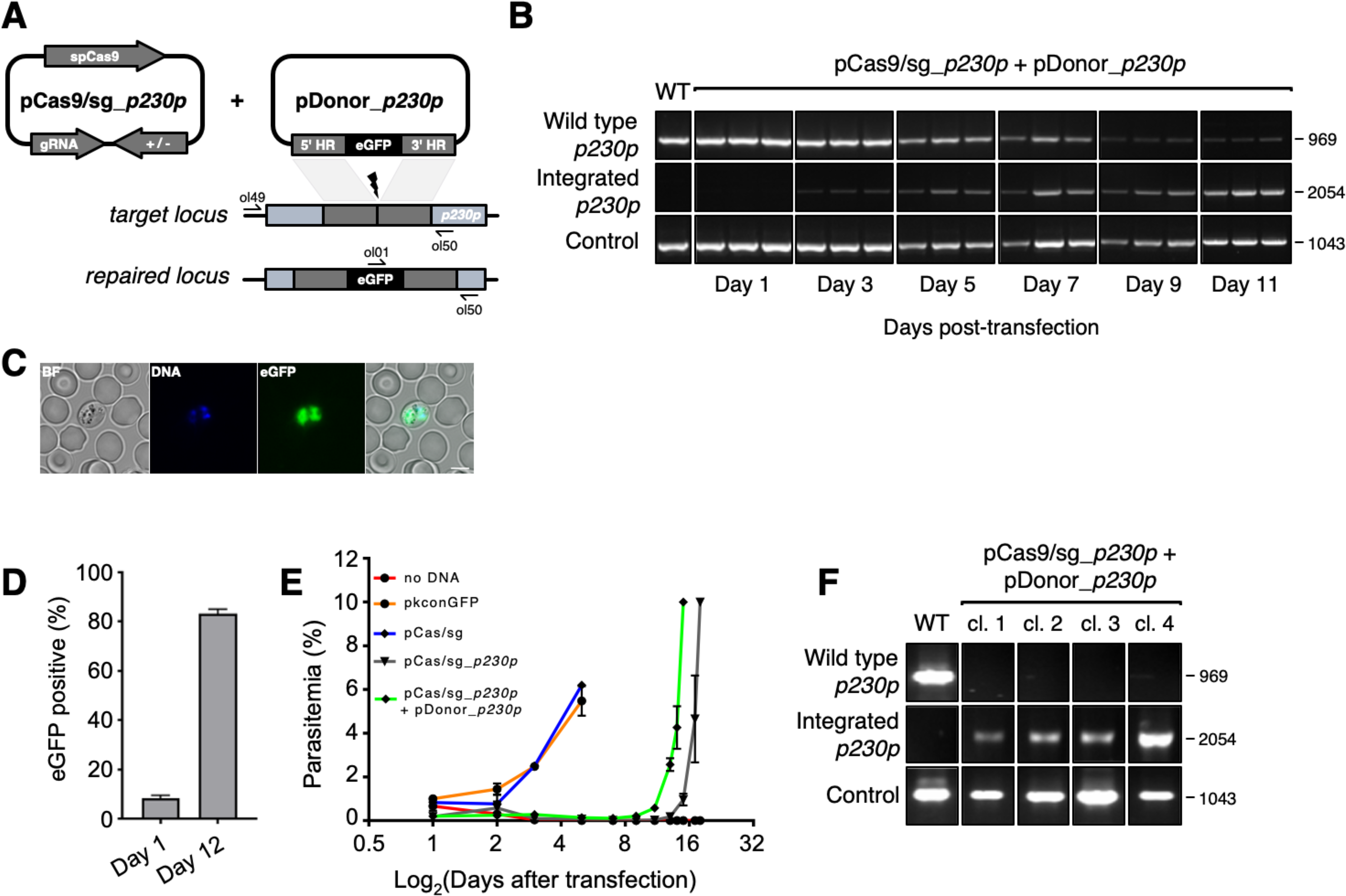
CRISPR-Cas9 genome editing in *P. knowlesi*. (A) Schematic of CRISPR-Cas9 strategy. Integration of the eGFP expression cassette into the target *p230p* locus via homologous recombination. Arrows indicating oligo positions for diagnostic PCRs. (B) Parasites transfected with pCas9/sg_*p230p* and pDonor_*p230p* plasmids were analysed with diagnostic PCRs on consecutive days after transfection. PCR reactions detecting the wild type locus (ol49+ol50), integration locus (ol01+ol50) and a control PCR targeting an unrelated locus (ol75+ol76) using approximately 3 ng/μl genomic DNA. For each day, three transfections are shown. (C) Representative live microscopy image of eGFP positive schizont transfected with pCas9/sg_*p230p* and pDonor_*p230p* plasmids. Scale bar represents 5 μm. (D) Proportion of eGFP positive parasites (%) counted after transfection with pCas9/sg_*p230p* and pDonor_*p230p* plasmids to show transfection efficiency on day 1 and integration efficiency after reaching culture reached 0.5 % parasitemia (day 12) (n=3). Error bars denote ± 1 SD. (E) Graph shows change in parasitemia (%) over time for parasite lines transfected with the dual plasmid Cas9 targeting vectors (pCas9/sg_*p230p* and pDonor_*p230p)*, controls without an sgRNA (pCas9/sg), without homology repair template DNA (pCas9/sg_p230p) or with no DNA. A fifth control reaction shows outgrowth of an episomal control plasmid (pkconGFP) (n=3). Parasites were placed under drug selection on day 1. Error bars denote ± 1 SD (F) Parasites transfected with pCas9/sg_*p230p* and pDonor_*p230p* plasmids were cloned by limiting dilution and four clones analysed by diagnostic PCR.

### A three-step PCR method enables rapid, cloning-free generation of donor constructs

*P. knowlesi* readily accepts linearised plasmids for homologous recombination (20, 25, 33), so we next tested whether we could use a PCR-based approach for scalable generation of repair templates. As no selectable marker is used within the repair template, this could be easily produced by using PCR to fuse 5’ and 3’ HRs with the region containing the desired insertion, dispensing with the need for a plasmid back-bone. Modifying a method used for homologous recombination in *P. berghei* (34), we developed a three-step PCR scheme which first amplified the eGFP cassette and 400 bp HRs with eGFP cassette adaptors separately, with the second and third reactions fusing each HR to the eGFP cassette in turn (Figure 2A). The addition of nested primers for the second and third PCR step removed background bands and improved robustness. The final PCR construct (HR1-eGFPcassette-HR2) was transfected along with the pCas9/sg_*p230p* plasmid, and resultant parasite lines demonstrated integration by PCR (Figure 2B), and an eGFP positivity rate of 74 % (± 8 SD), similar to that seen for the pDonor_*p230p* plasmid (Figure 2C). The use of a high-fidelity, high-processivity polymerase for the construct production allowed each reaction to be completed in 40-90 minutes, thus providing a rapid method for generating repair templates.

**Figure 2:**
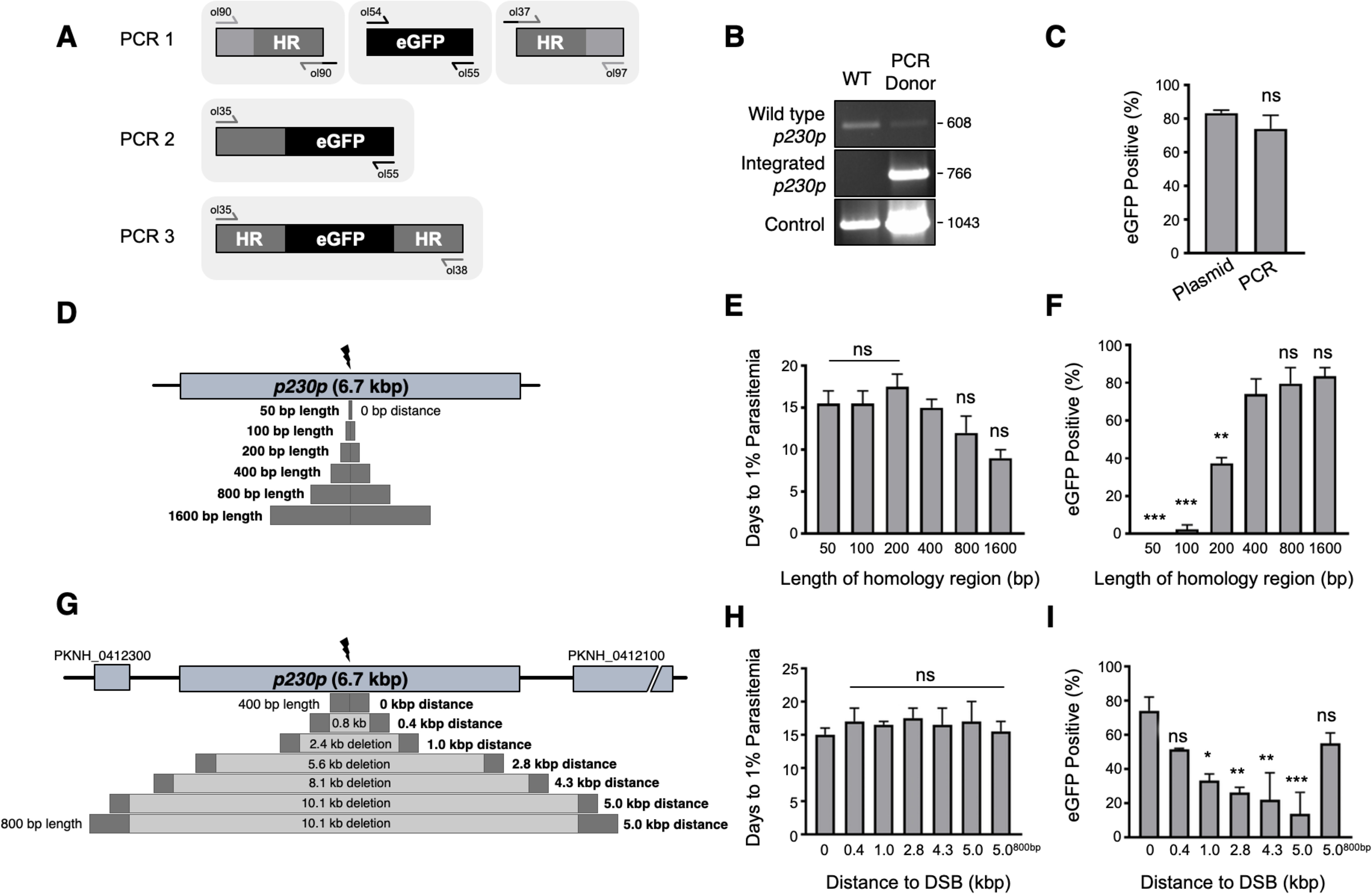
Fusion PCR based approach enables cloning-free production of homology repair templates and evaluation of key parameters for efficient homology-driven repair. (A) Schematic of the nested PCR method to generate linear donor constructs for transfection. First, homology regions (HRs) and eGFP cassette were amplified by PCR with 20 bp overhangs and gel extracted. In a second nested step HR1 and eGFP cassette were fused and again in the third step the HR1-eGFP product was fused with HR2. (B) Parasites transfected with pCas9/sg_*p230p* and PCR repair template (PCR donor), comprised of an eGFP cassette and 400bp HRs, were analysed with diagnostic PCRs amplifying the wild type p230p locus (ol49+ol50), integration locus (ol01+ol50) and a control targeting an unrelated locus (ol75+ol76). (C) After selection for integration, the proportion of eGFP positive parasites (%) was determined by fluorescent microscopy and compared between Cas9 transfections made with 400 bp HR plasmid (pDonor_*p230p*) or 400 bp HR PCR donor DNA. Data points represent the mean and error bars indicate ± 1 SD of two independent experiments (n=2). (D) The *p230p* locus was targeted using PCR donor DNA constructs using HRs with 50-1600 bp length. The bar chart shows, for each of the constructs with HRs of 50 to 1600 bp length, (E) the number of days for transfections to reach 1 % parasitemia and (F) proportion of eGFP positive parasites (%) after selection. All transfections were carried out in two independent experiments. (G) The *p230p* locus was targeted using PCR donor DNA constructs with HRs placed at varying distance from the Cas9 induced double strand break (DSB). For each construct based on distance to the DSB, the bar chart shows, (H) the number of days for transfections to reach 1 % parasitemia and (I) proportion of eGFP positive parasites (%) after selection. Data points represent the mean and error bars indicate ± 1 SD of two independent experiments (n=2). Results were all compared to the 400 bp HR construct at 0 kb from DSB using a one-way ANOVA with Dunnett’s multiple comparison of means. ns > 0.05, * < 0.05, ** < 0.01, *** < 0.001.

### Longer HRs increase the integration efficiency and offsets DSB distance efficiency loss

We next used this PCR approach to investigate the optimal parameters and limits of the Cas9 system in *P. knowlesi*. Varying the length of HRs targeting the same *p230p* locus (Figure 2D), allowed us to determine the effect on integration efficiency as well as the size limits of the PCR approach. The largest construct generated in this way was 6.1 kb in length (2x 1.6 kb HRs flanking the 2.9 kb eGFP expression cassette). Attempts to generate a larger 9.3 kb construct (2x 3.2 kb HRs) failed during the final PCR step. PCR yields were lower for larger constructs, with the 6.1 kb construct yielding half that of the 3.7 kb construct. PCR repair templates with HRs ranging from 50-1600 bp generated single specific bands with exception of the 400 bp HRs which contained an additional lower band, due to a primer additionally annealing to a repeat region in HR1 (S2B Figure). The PCR constructs were transfected together with the pCas/sg_*p230p* plasmid and integration efficiency monitored (Figure S2A). All 6 HRs lengths produced evidence of integration by PCR, but the efficiency rapidly declined with shorter HR length (S2D Figure). Parasites transfected with 800 and 1600 bp HR constructs were the fastest to reach 1 % parasitemia on day 12 and 9 post transfection, respectively (Figure 2E). For the 50 and 100 bp HR constructs no eGFP positive parasites were detected by fluorescence microscopy suggesting very low targeting efficiencies. Constructs with HRs >400 bp provided GFP positivity, ranging from 79 and 81 % (Figure 2F), which taken together with PCR yields and transfection recovery time suggest an optimal HR length of at least ~800 bp.

To undertake large gene deletion or replacement experiments, HRs may need to be placed at a distance from the Cas9-induced DSB, and it is well known in other systems that efficiency rapidly declines with distance to DSB (35). In *P. falciparum* integration efficiencies decrease drastically with distance over 250 bp from the PAM site (36). To determine how distance from DSB affected efficiency of integration, we used the same *p230p* PAM site and moved our 400 bp HRs varying distances away from the DSB, ranging from 0 - 5 kb (Figure 2G and S2D Figure). Whilst all transfections were PCR positive for integration and reached 1 % parasitemia at similar times (14-20 days) (S2D Figure and Figure 2H), the integration efficiency declined with distance from DSB. This decline was surprisingly small, with HRs placed even 5 kb away from either side of the DSB yielding a 14 % (± 18 SD) integration efficiency (Figure 2I). Interestingly, we found that extending HR length to 800 bp restored integration efficiencies to 54.8 % (± 8.7 SD) at a 5 kb distance from DSB (Figure 2I). Thus, HR length can directly offset efficiency losses due to distance from DSB and this system can readily remove genes at least as large as 10 kb in size from a single PAM site, accounting for ~98 % of genes in the *P. knowlesi* genome (37).

### Cas9-based PCR constructs enable rapid and flexible gene tagging in *P. knowlesi*

Having demonstrated consistent performance of an sgRNA sequence in the sgRNA/Cas9 suicide vector and PCR constructs for targeting a single control locus, we next sought to determine how robust the system is for targeting a range of loci. We therefore used the PCR-based approach for fusion of fluorescent or epitope tags to proteins of interest (Figure 3A). For C-terminal tags, the PCR repair templates were generated by creating fusions of the tag with HRs targeting the 3’end of the gene and the 3’UTR. Similarly, N-terminal tag repair templates were created by flanking the tag with HRs targeting the 5’UTR and 5’end of the coding region. In each case a PAM site was selected that crossed the stop codon (for C-terminal) or start codon (for N-terminal) such that integration of the tag alone, with no other exogenous sequence, was sufficient to disrupt the PAM site. For genes, such as the Chloroquine Resistance Transporter (CRT), where the PAM site preceded the stop codon, intervening sequences were recodonised when generating the 5’HR to disrupt the PAM site using silent mutations. We selected five genes with disparate subcellular locations and functions to test this approach: the micronemal protein apical membrane antigen 1 (AMA1) (38), rhoptry neck protein 2 (RON2) (39), inner membrane complex protein myosin A (MyoA) (40), digestive vacuole membrane protein involved in drug resistance CRT (41), and a protein involved in artemisinin resistance in cytoplasmic foci Kelch13 (K13) (42). A single sgRNA was selected for each, and repair templates were generated by fusion PCR to incorporate an eGFP, mCherry (both with 24 bp glycine linker) or a hemagglutinin (HA) tag (S3A-E Figure). An N-terminal tag was used for K13, as previous work in *P. falciparum* suggested that C-terminal tagging affected parasite growth (42), and C-terminal tags used for all the other targets. All lines grew up quickly after transfection, reaching 1 % after between 8 and 15 days, and PCR analysis indicated that correct integration had occurred (Figure 3B). Whilst it is, to our knowledge, the first time each of these proteins have been tagged in *P. knowlesi*, all demonstrated localisation patterns were consistent with previous reports for *P. falciparum* (Figure 3C). AMA1, MyoA and K13 showed clear bands at the expected size on western blots. The CRT-eGFP fusion protein showed a band at ~50 kDa, in line with work in *P. falciparum* which showed CRT-eGFP migrates faster than its predicted size of 76 kDa (S3F Figure) (41). We were unable to visualise a band for RON2-HA most likely due to poor blotting transfer of this 240 kDa protein. Together, these results demonstrate that the fusion PCR approach can be used to tag *P. knowlesi* genes rapidly and robustly at a variety of loci. Analysis of equivalent *P. falciparum* loci revealed only 2/5 had suitably positioned PAM sites, and equivalent UTR regions had an average GC-content of only 11.8 % (36 % for *P. knowlesi*), suggesting a similar approach would have been more challenging in *P. falciparum* (Table S1).

**Figure 3:**
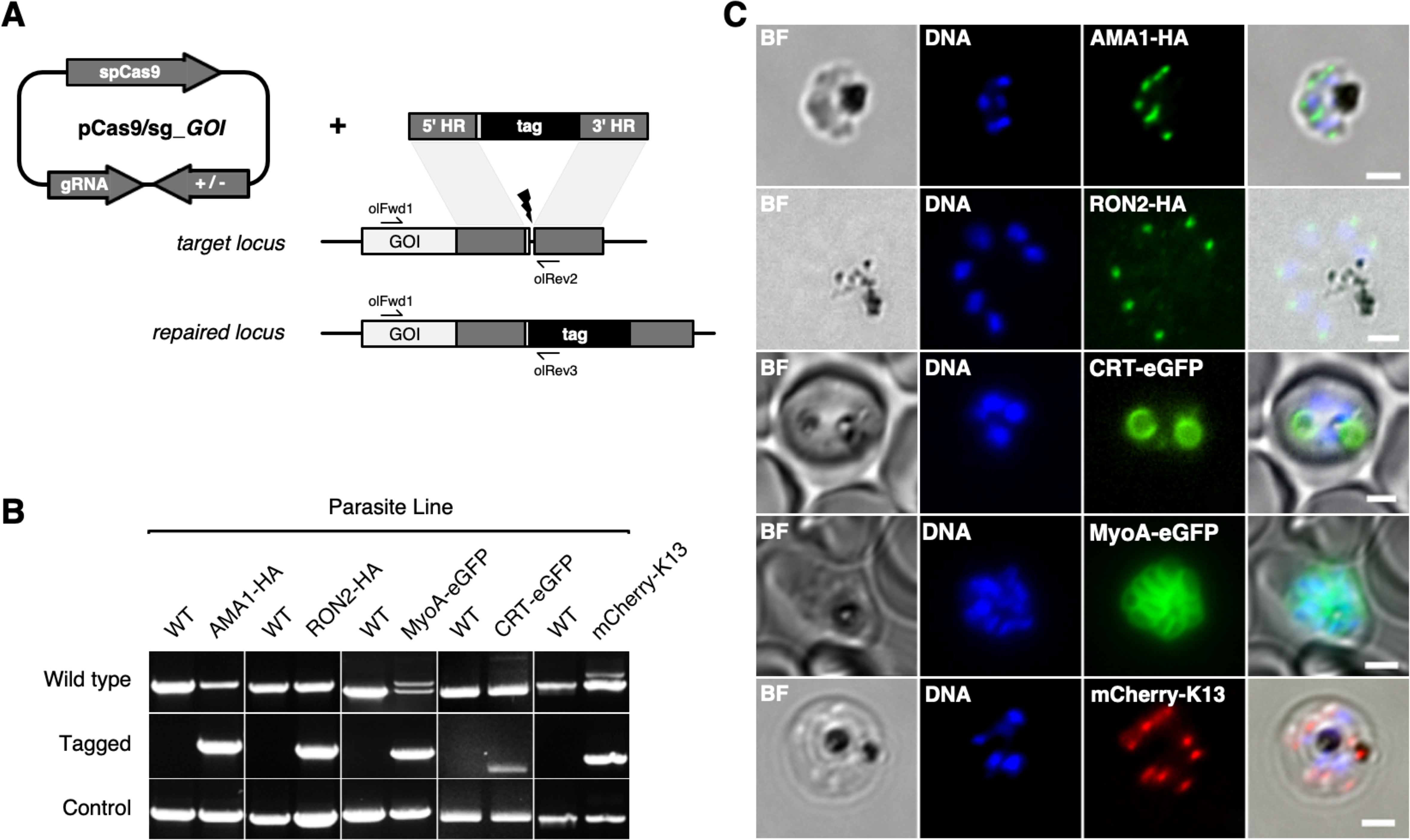
CRISPR Cas9 PCR repair templates enable rapid and flexible tagging of parasite proteins. (A) Schematic of CRISPR-Cas9 system for tagging. pCas9/sg plasmid with gene of interest (GOI) specific sgRNA, is combined with repair template generated by fusion PCR. Lightning bolt indicates Cas9 induced double strand break, which is repaired by insertion of the desired tag. (B) Diagnostic PCRs specific to each GOI locus were carried out to amplify the wild type locus (schematic positions olFwd1 +olRev2), integration locus (schematic positions olFwd1 +olRev3) and a control targeting an unrelated locus (ol75+ol76). List of specific primers used for each GOI is shown in Table S2. As no DNA is removed in this process, the wild type specific locus primers also generate slightly larger amplicons in tagged lines, which can be seen as double bands for both the Myosin A and K13 PCRs.(C) Representative immunofluorescence images of HA-tagged Apical membrane antigen-1 (AMA1-HA) and Rhoptry neck protein 2 (RON2-HA) parasite lines, and live cell imaging of Chloroquine Resistance Transporter-eGFP (CRT-eGFP), Myosin A-eGFP (MyoA-eGFP) and mCherry-Kelch13 (K13). Panels shows brightfield (BF), DNA stain (blue) and anti-tag antibodies/live fluorescence (green or red) of schizonts stage parasites from each line. Scale bars represent 2 µm.

### Transgenic *P. knowlesi* orthologue replacement lines provide surrogates for *P. vivax* vaccine development and DBP tropism studies

Having demonstrated the utility of this technique for rapidly manipulating genes of interest, we next sought to use this system to study *P. vivax* biology. The orthologous RBC ligands PkDBPα and PvDBP, mediate host cell invasion by binding to the DARC receptor on human RBCs in *P. knowlesi* and *P. vivax*, respectively (5–8). PvDBP is currently the lead vaccine candidate for a *P. vivax* blood stage vaccine (12–14), thus *P. knowlesi* could provide an ideal surrogate for vaccine testing in the absence of a robust in vitro culture system for *P. vivax*. Whilst likely functionally equivalent, the DBP orthologues are antigenically distinct (~70 % amino acid identity in binding region II) so we used genome-editing tools to generate transgenic *P. knowlesi* parasites in which DARC binding is provided solely by PvDBP. We first carried out an orthologue replacement (OR) of the full-length PkDBPα with PvDBP in the *P. knowlesi* A1-H.1 line (PvDBP^OR^) – using a recodonised synthetic PvDBP gene flanked by HRs targeting the 5’ and 3’UTRs of the PkDBPα gene (S4A Figure). Once integrated, this deletes the PkDBPα gene and places the PvDBP gene under control of the PkDBPα regulatory sequences, enabling a precisely matched expression profile. As a control we also exchanged PkDBPα with a recodonised PkDBPα gene (PkDBPα^OR^) using the same sgRNA (S4C Figure). Successful integration was readily achieved and limiting dilution cloning resulted in 100 % integrated clones for PkDBPα^OR^ and 40 % for PvDBP^OR^ (Figure 4A). The PkA1-H.1 line relies on the DARC receptor for invasion of human RBCs (20) and PkDBPα is required to mediate this interaction (7), thus the successful replacement indicates that the Pv orthologue can fully complement its role in DARC binding and parasite invasion.

**Figure 4:**
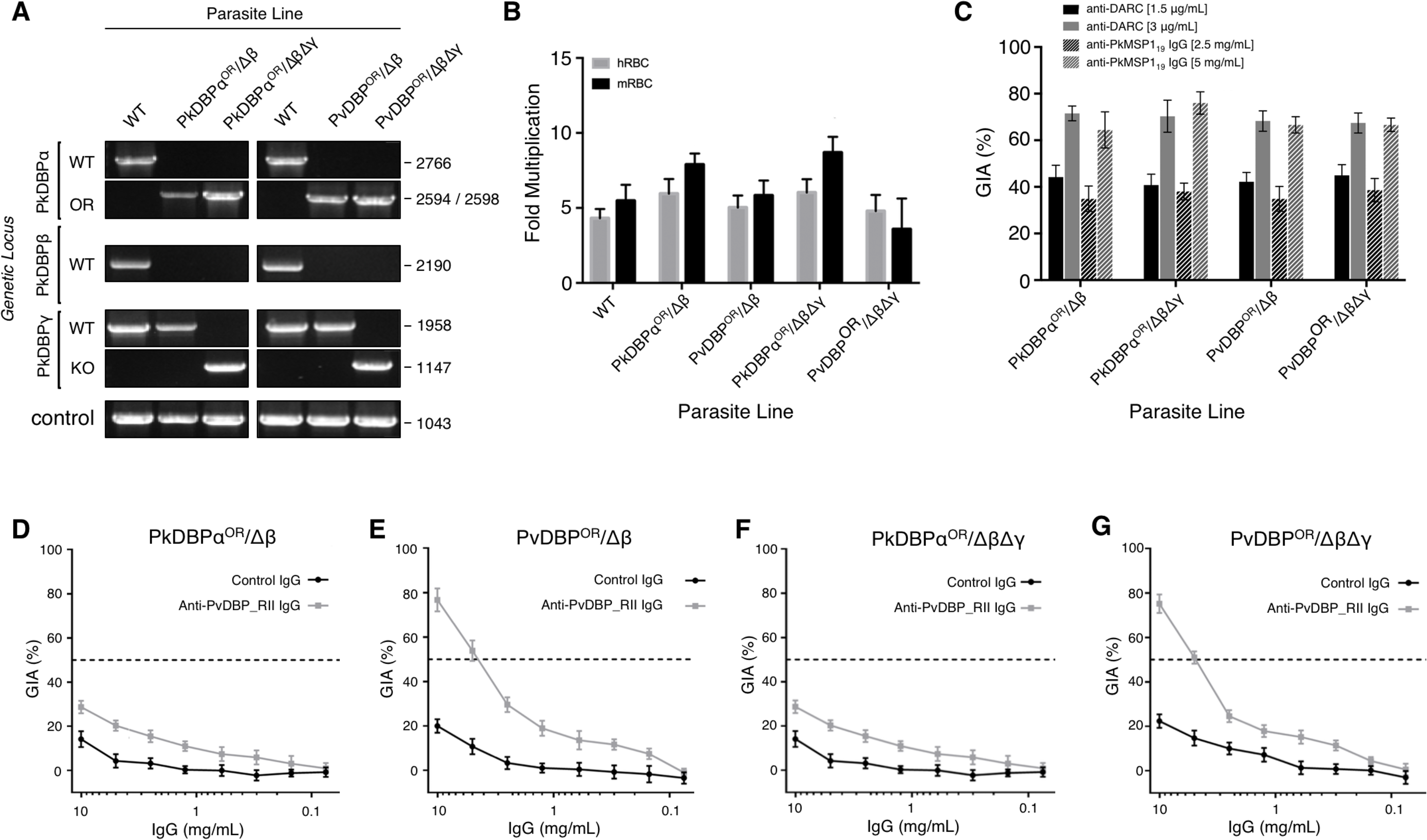
Transgenic *P. knowlesi* orthologue replacement lines provide surrogates for *P. vivax* vaccine development. (A) The *P. knowlesi* Duffy binding protein α (DBPα) gene was targeted for replacement with either a recodonised PkDBPα or *P. vivax* DBP repair template. Sequencing revealed a loss of ~44 kb in chromosome 14, that includes loss of PkDBPβ (PkDBPα^OR^/Δβ and PvDBP^OR^/Δβ). These lines were then subsequently modified to knockout PkDBPγ (PkDBPα^OR^/ΔβΔγ and PvDBP^OR^/ΔβΔγ). Parasite lines were analysed using PCR reactions detecting the wild type (WT) locus PkDBPα (ol186+ol188), orthologue replacement (OR) locus of PkDBPα^OR^ (ol186+ol189) or PvDBP^OR^ (ol186+ol187), WT PkDBPβ locus (ol480+481), WT locus of PkDBPγ (ol483+ol484), KO locus of PkDBPγ (ol483+ol258) and a control PCR targeting an unrelated locus (ol75+ol76). (B) Bar chart showing mean fold replication of parasites lines in a FACS-based invasion assays over one growth cycle (24 h). Assays were carried out in eight independent experiments for human blood (hRBC) and three independent experiments for *Macaca fascicularis* blood (mRBC). Error bars indicate ± 1 SD. Data points represent the mean single cycle growth rate. Replication rates of the parasites lines were compared by using unpaired t-tests comparing means. There are significant differences in fold multiplication rates of PkDBPα^OR^/Δβ against PvDBP^OR^/Δβ in mRBCs (p<0.05), and PkDBPα^OR^/ΔβΔγ against PvDBP^OR^/ΔβΔγ in hRBCs (p<0.05) and mRBCs (p<0.05). (C) Graph showing growth inhibition activity (GIA, %) of anti-DARC nanobody at 1.5 and 3 μg/ml and anti-MSP1_19_ purified total rabbit IgG at 2.5 and 5 mg/ml on the parasite lines. Data points represent the mean and error bars indicated ± 1 SD of triplicate test wells (n=3). (D-G) Graphs showing % GIA of anti-PvDBP-RII IgG purified from rabbit serum against transgenic *P. knowlesi* lines. Each panel shows the % GIA of a dilution series of total IgG purified from sera of PvDBP_RII (SalI)-immunized rabbits as well as control IgG from the pre-immunisation sera of the same rabbits on (D) PkDBPα^OR^/ Δβ (E) PvDBP^OR^/ Δβ, (F) PkDBPα^OR^/ ΔβΔγ line, (G) PvDBP^OR^/ ΔβΔγ line. Data points represent the mean and error bars indicate ± 1 SD of 6 replicates.

*P. knowlesi* contains two DBPα paralogues, DBPβ and DBPγ, which are highly homologous at the nucleotide (91-93 % identity) and amino acid (68-88 % identity) levels, but are thought to bind to distinct sialic acid-modified receptors unique to macaque RBCs (16). The PkDBPα sgRNA was carefully designed to be distinct to equivalent DBPβ and DBPγ target sequences (85 % identical to DBPγ and 47.8 % to DBPβ), because, as in other systems, off-target Cas9-induced DSBs are a major issue (43, 44). We therefore sequenced the four most similar target sequences, including one in DBPγ, in the PvDBP^OR^ lines (Table S2) and did not detect any off-target mutations, suggesting that as for other malaria parasites (22) the absence of non-homologous end joining (26) ameliorates the potential for off-target mutations. However, diagnostic PCRs for DBPβ failed, as well as PCRs in genes flanking the DBPβ locus. Whole genome sequencing revealed that in one of two independent PkDBPα^OR^ and PvDBP^OR^ clones of ~44 kb truncation at one end of chromosome 14 had occurred (S5 Figure), which also harbours DBPβ, resulting in lines PkDBPα^OR^/Δβ and PvDBP^OR^/Δβ. The loss of the ~44 kb of chromosome 14 is also present in parasites that have been transfected simultaneously with pCas/sg_*p230p*, suggesting that the 44 kb deletion occurred in the A1-H.1 parental parasite line prior to transfection and was not an artefact caused by targeting DBPα. Similar spontaneous deletions have been reported previously, including ~ 66 kb loss at the other end of chromosome 14 in the *P. knowlesi* A1-C line maintained in cynomolgus macaque blood that included the invasion ligand NBPXa (33), and a deletion of DBPγ in the PkYH1 line at the end of chromosome 13 (16). Furthermore, the PAM site of the DBPα targeting guide sequence is absent in DBPβ (S4C Figure) which makes it unlikely that the disruption of DBPβ was induced by Cas9 during DBPα targeting.

Having established accurate targeting of the PkDBPα locus, we investigated the role of the paralogues in human and macaque red cell invasion and whether they could interfere with inhibitory effects of test antibodies. We took advantage of the line with the spontaneous DBPβ loss and then used pCas/sg plasmid recycling to additionally delete the DBPγ locus, generating PkDBPα^OR^/ΔβΔγ and PvDBP^OR^/ΔβΔγ. The final PvDBP^OR^/ΔβΔγ clonal line was subjected to whole genome sequencing to verify changes at all three loci, and this confirmed precise targeting of the PvDBP allele swap into the PkDBPα locus, and complete deletion of the DBPγ open reading frame in the PkDBP γ locus (S5 Figure).

Analysis of the wild type line, and the four transgenic lines (PkDBPα^OR^/Δβ, PvDBP^OR^/Δβ, PkDBPα^OR^/ΔβΔγ and PvDBP^OR^/ΔβΔγ) revealed no difference in growth rate in human RBCs (Figure 4B), confirming that the *P. vivax* protein was able to fully complement the role of its *P. knowlesi* orthologue, and PkDBPβ and γ proteins are dispensable for invasion of humans RBCs (16). To investigate how these modifications affect host tropism we compared the growth rates in human RBCs with growth rates in RBCs from the natural host of *P. knowlesi*, the long tailed macaque (*Macaca fascicularis*). Even human culture-adapted *P. knowlesi* retains a strong preference for macaque cells (20, 21, 33) and it has been hypothesized that the additional invasion pathways provided by DBPβ and DBPγ are in part responsible for the increased invasion efficiency in macaque RBCs (16). Interestingly, loss of DBPβ and DBPγ in the PkDBPα^OR^/ΔβΔγ line did reduce parasite replication in macaque RBC (Figure 4B), with the lines retaining a macaque preference ratio (macaque fold growth/human fold growth) of 1.43, similar to both the wild type (1.26) and PkDBPα^OR^/Δβ (1.33). This demonstrates that both proteins are dispensable for invasion of macaque RBCs, and PkDBPα alone is sufficient to retain full invasion efficiency in macaque cells. Unlike *P. knowlesi*, *P. vivax* is unable to infect macaques, and sequence differences between the DARC receptor in the two hosts have been suggested to underlie this restriction (11). Whilst macaque invasion rates and host preference ratio (1.16) were unaffected for the PvDBP^OR^/Δβ line, the additional deletion of DBPγ in the PvDBP^OR^/ΔβΔγ line resulted in a 40% reduction in macaque invasion rates (Figure 4B) which caused a shift to human RBC preference with a ratio of 0.75. This suggests that in the absence of redundant DBP pathways, PvDBP is less effective at facilitating invasion of macaque cells than of human cells, but nevertheless can support invasion of both host cell types.

Growth inhibition activity (GIA) assays revealed that all lines remained equally susceptible to invasion inhibition by both an anti-DARC camelid nanobody CA111 (45) and a polyclonal αPkMSP1_19_ antibody (Figure 4C). In contrast, purified IgGs from polyclonal rabbit sera raised against PvDBP-RII, demonstrated low-level GIA activity for wild type and PkDBPa^OR^/ΔβΔγ lines (~30 % inhibition at 10 mg/ml) but a significantly stronger GIA activity against the PvDBP^OR^ and PvDBP^OR^/ΔβΔγ lines, reaching a maximum inhibition of ~75 % at 10 mg/ml and around 50 % at 4 mg/ml (Figure 4 D-G). The PvDBP^OR^ parasite lines could thus be readily inhibited by antibodies against the *P. vivax* protein and the PkDBPα orthologues γ and β appeared to play no interactive role. We thus have created a transgenic *P. knowlesi* model, modified at two separate loci which recapitulates the *P. vivax* DBP invasion pathway. This parasite line is a vital new tool in PvDBP vaccine development.

## Discussion

In this work, we adapt CRISPR-Cas9 genome editing to the zoonotic malaria parasite *P. knowlesi*. Whilst various approaches for CRISPR-Cas9 have been used for other malaria parasites (22, 29, 44, 46), here we combine a plasmid containing a single recyclable positive selection marker with a fusion PCR-based approach for generation of repair templates. This allows for seamless insertion or deletion at any location within a gene and unlimited iterative modifications of the genome. Genome-wide reverse genetics screens have been applied with great success to the rodent malaria parasite, *P. berghei* (47, 48), but they have remained challenging for *P. falciparum*, and impossible for *P. vivax*. The tools presented here will enable scalable construct assembly and genome-wide systematic knockout or tagging screens in an alternative human infective species, thus providing a complementary tool to address both shared and species-specific biology. The analysis of lines with multiple tagged or deleted genes is particularly valuable for multigene families with highly redundant functions, as exemplified by our modification of all three *P. knowlesi* DBP genes.

Here we investigate key parameters associated with successful genome editing and show that the process is also highly robust; targeting of the *p230p* locus demonstrated successful editing for 25/25 transfections and only 1/10 sgRNAs targeting different loci failed to generate an edited line. The failure of an sgRNA guide (AGAAAATAGTGAAAACCCAT) designed to target the DBPβ locus, a non-essential gene, suggests that multiple guides may need to be tested for some loci. We did not detect any off-target effects, consistent with other reports of CRISPR-Cas9 use in malaria parasites (22, 29, 44, 46). Negative selection of the pCas9/sg plasmid then enables generation of markerless lines allowing unlimited iterative modifications of the genome, with each round requiring only ~30 days (including dilution cloning). We systematically tested key parameters associated with successful genome editing and found increasing HR length enhanced integration efficiency proportionately, a trend seen in both *P. falciparum* and *P. berghei* (31, 44, 49). Whilst integration was detected with HRs as short as 50 bp, efficient editing was achieved with HRs between 200-800 bp. We were also able to examine how distance from the DSB affected editing efficiency. Whilst in other systems editing efficiency decreases rapidly as the DSB distance increases, we saw only a steady decline with distance, an effect which could be ameliorated by simply increasing HR length.

By applying these techniques to the *P. knowlesi* and *P. vivax* DBP family we have been able to examine the role of these genes in host and reticulocyte tropisms of the two species. Even after long-term adaptation to culture with human RBCs, *P. knowlesi* parasites can retain a strong preference for invasion of macaque RBCs reticulocytes (16, 33). Both DBPγ and DBPβ have been shown to bind to proteins with a distinct sialic acid residue found in non-human primates, but absent in humans (16). Deletion of these genes had no effect on invasion of human RBCs, but interestingly also had no effect on invasion efficiency in macaque RBCs either, demonstrating that PkDBPα alone is sufficient to retain full invasive capacity and that the DBP proteins are not responsible for the macaque cell preference retained in the A1-H.1 human adapted line. Despite being closely related to *P. knowlesi* and other macaque infecting species such as *P. cynomolgi*, *P. vivax* cannot infect macaques and the PvDBP protein has been suggested to play a role in enforcing this tropism, as key interacting residues are missing within the macaque DARC protein (11). *P. knowlesi* parasites expressing PvDBP in the absence of DBP paralogues demonstrate a clear reduction in invasion capacity in macaque cells, resulting in an overall shift towards preference for human cells consistent with a PvDBP binding macaque DARC less efficiently. Nevertheless as invasion capacity remained quite close to that seen for human RBCs it seems unlikely that the PvDBP protein alone represents a significant barrier to *P. vivax* infection of macaques. Another key difference between the two species is that unlike *P. knowlesi, P. vivax* has a strict restriction to invasion of reticulocytes. A second family of RBC binding proteins, known as the reticulocyte binding-like proteins (RBPs) have previously been implicated in this tropism. More recently, the PvDBP protein itself has been implicated with work using recombinant PvDBP-RII suggesting that whilst DARC is present on both reticulocytes and mature normocytes, changes during red cell maturation mean that DARC is only accessible to PvDBP binding in young reticulocytes (10). Here we show that transgenic *P. knowlesi* parasites using PvDBP for invasion have no such restriction, invading human RBCs (which typically contain less than 0.5% reticulocytes) with the same efficiency as those expressing PkDBP – thus providing compelling evidence that PvDBP plays no role in the reticulocyte tropism. Further, recent work determining that PvRBP2b, which lacks an orthologue in *P. knowlesi*, binds to the reticulocyte specific marker CD71 (50) further asserts the RBPs as the key to reticulocyte tropism. Importantly, the ability to compare and contrast activity of Pk/Pv DBP family members in parasitological assays will provide a vital new tool to test hypotheses and models arising from studies that have until now relied on assays using recombinant protein fragments.

Efforts to develop a *P. vivax* vaccine to elicit antibodies against the lead candidate PvDBP have predominantly relied on using ELISA-based assays, which assess the ability of antibodies to block recombinant PvDBP-RII binding to DARC (18), but are likely to be less informative than parasitological assays. Some epitopes identified in recombinant protein assays may be inaccessible in the context of invasion and it is also possible that not all inhibitory antibodies directly block receptor engagement. DARC-DBP binding is only one step in the multi-step invasion process, with subsequent conformational changes and potential downstream signalling roles for the protein (51). The full-length DBP antigen is 140 kDa which contains a C-terminal transmembrane domain and as such structural and biochemical analysis of the protein has almost exclusively focused on the PvDBP-RII fragment alone. The *P. knowlesi* PvDBP^OR^ line thus provides an opportunity to interrogate the function of the full-length protein. Whilst efforts to standardise ex vivo *P. vivax* assays have been successful (15), they remain hugely challenging, low throughput and rely on genetically diverse *P. vivax* clinical isolates, that are maintained in culture for a only a single cycle of RBC invasion. A vaccine against *P. vivax* must ultimately elicit antibodies with strain-transcending inhibitory activity, but the ability to test on a defined genetic background can provide a significant advantage when it comes to benchmarking and prioritising target epitopes and characterising sera raised against them. Here we use the PvDBP sequence from the SalI reference strain, but multiple lines expressing distinct PvDBP variants could be generated in future, to systematically examine inhibition in heterologous strains. Isolates refractory to a given test antibody in ex vivo assays can then be sequenced and direct the generation of new transgenic *P. knowlesi* PvDBP^OR^ variant lines to support rational vaccine development. These assays in turn can provide vital triaging for non-human primate models, and controlled human challenge infections (13, 15, 17, 19) – both of which carry the imperative to ensure that only highly optimised antigens are tested.

The transgenic *P. knowlesi* OR lines developed here represent the ideal platform for scalable testing of polyclonal and monoclonal sera from vaccine trials and natural *P. vivax* infections. This will enable detailed investigation of epitopes providing invasion inhibitory activity and a means for systematic development of a strain-transcending vaccine. Our work also revealed low-level cross-reactivity of PvDBP-RII antibodies against *P. knowlesi* and suggests cross-immunity between the two species could exist in the field, which may have a significant impact on disease outcome. Understanding the precise epitopes involved could facilitate development of a dual species vaccine, and epitopes conserved across species are also more likely to be conserved across polymorphic strains of *P. vivax*. The same approach could readily be applied to other potential vaccine candidates, novel drug targets or to investigate mechanisms of drug resistance, which are also thought to differ between *P. falciparum* and *P. vivax* (52).

In conclusion, we demonstrate that adaptation of CRISPR-Cas9 genome editing to *P. knowlesi* provides a powerful system for scalable genome editing of malaria parasites and can provide critical new tools for studying both shared and species-specific biology.

## Methods

### Macaque and Human RBCs

*Macaca fascicularis* blood was collected by venous puncture. Animal work was reviewed and approved by the local National Institute for Biological Standards and Control Animal Welfare and Ethical Review Body (the Institutional Review Board) and by the United Kingdom Home Office as governed by United Kingdom law under the Animals (Scientific Procedures) Act 1986. Animals were handled in strict accordance with the “Code of Practice Part 1 for the housing and care of animals (21/03/05)” available at https://www.gov.uk/research-and-testing-using-animals. The work also met the National Centre for the Replacement Refinement and Reduction of Animals in Research (NC3Rs) guidelines on primate accommodation, care, and use (https://www.nc3rs.org.uk/non-human-primate-accommodation-care-and-use), which exceed the legal minimum standards required by the United Kingdom Animals (Scientific Procedures) Act 1986, associated Codes of Practice, and the US Institute for Laboratory Animal Research Guide. Human blood (Duffy (FY) positive) was obtained from the United Kingdom National Blood Transfusion Service under a research agreement.

### Parasite Maintenance, transfection and dilution cloning

Parasites were maintained in complete media, comprising RPMI 1640 (Invitrogen) with the following additions: 2.3 g/L sodium bicarbonate, 4 g/L dextrose, 5.957 g/L HEPES, 0.05 g/L hypoxanthine, 5 g /L Albumax II, 0.025 g/L gentamycin sulfate, 0.292 g/L L-glutamine, and 10 % (vol/vol) horse serum as described previously (33). Parasites were synchronized by using gradient centrifugation with 55 % nycodenz (Progen) in RPMI to enrich schizonts, followed by a two-hour incubation with 4-[7-[(dimethylamino)methyl]-2-(4-fluorphenyl)imidazo[1,2-*a*]pyridin-3-yl]pyrimidin-2-amine (compound 2) which inhibits parasite egress(31). Incubations in compound 2 longer than 2 hours led to degeneration of schizonts and reduction in invasive capacity.

Tightly synchronized mature schizonts were transfected as described previously using the Amaxa 4D electroporator (Lonza) and the P3 Primary cell 4D Nucleofector X Kit L (Lonza)(20). 10 μl DNA including at 20 µg repair template pDonor_*p230p* and 20 μg pCas9/sg_*p230p* plasmid was used for transfections to generate eGFP expressing lines. 10 μl DNA including 15 μg repair template and 7 μg pCas9/sg_*p230p* plasmid was used for transfections to integrate the eGFP expression cassette into the *p230p* locus with PCR repair templates. For generating tagged lines 10 μg pCas/sg_GOI plasmid and 20 μg PCR repair templates were used. After 24 h, and at daily intervals for 5 days, the medium was replaced with fresh medium containing 100 nM pyrimethamine (Sigma). Parasites were cloned out by limiting dilution. Parasites were diluted to 0.3 parasites/100 μl and 100 μl of 2 % haematocrit culture was transferred to 96 flat-bottom plates in culture medium containing 200 mM L-glutamax (Sigma). After 7 days the media was changed and 0.2 % fresh blood added. On day 11 the plate was screened for plaques, in an assay modified from *P. falciparum* (32). Plaque positive cultures were transferred to 24 well plates containing 1 ml media with 2 % haematocrit and used for genotyping.

### DNA Constructs and PCRs

Preparative DNA for plasmid cloning and PCR fusion constructs was amplified with CloneAmp (Takara) using the following cycle conditions: 32 cycles of 5 s at 98°C, 20 s at 55°C, and 5 s/kb at 72°C. Genomic DNA was prepared using DNeasy blood and tissue kit (Qiagen).

#### Cloning of pkcon_mCherry plasmid

The plasmid PkconGFP (20) was modified to replace the GFP coding sequence with mCherry using XmaI and SacII restriction sites. The mCherry sequence was amplified with primers fwd-ATAT<u>CCCGGG</u>ATGGTGAGCAAGGGCGAGGAG and rev-ATAT<u>CCGCGG</u>TTACTTGTACAGCTCGTCCATGCC.

#### Cloning of pCas/sg

The pUF1 plasmid (22) was modified by replacing the yDHODH expression cassette with hDHFR-yFCU fusion with PkEF1a 5’UTR and Pbdhfr 3’UTR using EcoRI and SacII. The PfU6 promoter for gRNA expression of the pL6 plasmid (22) was replaced with the PkU6 5’ regulatable region of 1244 bp (amplified with primers fwd-ATATCCATGGGGCCAGGGAAGAACGGTTAGAG and rev-atattcgcgagcgatgagttcctaggAATAATATACTGTAAC) using NruI and NcoI and the entire cassette inserted into the pCas9 plasmid with PvuI and ApaI restriction sites. Each target specific 20 bp guide sequence was chosen with the Protospacer software (http://www.protospacer.com/) (49), with off-target score < 0.03. On-target scores were retrieved from Benchling Software (53). All guide sequences are listed in Table S3. Subsequently each guide was inserted into the BtgZI linearized pCas/sg plasmid by In-Fusion cloning (Takara) using primers fwd-TTACAGTATATTATT(N20)GTTTTAGAGCTAGAA and rev-TTCTAGCTCTAAAAC(N20)AATAATATACTGTAA. Briefly, 50 bp primers pairs containing the 20 bp guide sequence flanked by 15 bp overhangs homologous to the 5’ and 3’ ends of pCas/sg were denatured by incubation at 95 °C for 10 min and annealed by slow cooling. 0.5 μM annealed primers and 50 ng BtgZI linearized pCas/sg vector were incubated with In-fusion Premix (Takarta) at 50°C for 15 min. The resulting plasmid was transformed into XL10 gold competent *E.coli* cells (Agilent). Plasmids for transfection were prepared by Midi-preps (QIAGEN) and ethanol precipitated. The DNA pellet was washed twice with 70 % ethanol and resuspended in sterile TE buffer.

#### Cloning of pDonor_*p230p*

a plasmid containing a multiple cloning site with SacII, SpeI, NotI, BlpI and NcoI was designed (subsequently called pDonor) and obtained from Geneart (Thermo Fisher Scientific). Homology region 1 (HR1) was amplified from A1-H.1 wild type genomic DNA with primers olFM007 and olFM008 and added with SacII and SpeI restriction sites. HR2 was amplified with primers olFM005 and olFM006 and was added with BlpI and NcoI sites. The eGFP cassette was amplified from the pPkconGFP*p230p* plasmid with primers olFM151and olFM152 and inserted into the plasmid with SpeI and BlpI sites. The final vector was linearised with PvuI restriction enzyme and ethanol precipitated as described above.

#### Cloning of pDonor_*pkdbpα*

Plasmid *pDonor* was modified by restriction cloning to include two 500 bp HRs from PkDBPa 5’ and 3’UTRs using primers olFM062 and olFM063 (adding SacII /SpeI sites) and primers olFM064 and olFM065 (adding NotI/NcoI sites) respectively. Recodonised sequences of PkDBPα and PvDBP of the *P. vivax* Salvador I strain flanked with SpeI and NcoI restriction sites were obtained from Geneart (Thermo Fisher Scientific) and subsequently cloned between both HRs of the modified pDonor plasmid using SpeI and NcoI sites. The resulting plasmid was linearised with PvuI restriction enzyme and ethanol precipitated as described above.

#### Cloning of pDonor_ *pkdbp*γ

Plasmid *pDonor* was modified by restriction cloning to include two HRs from PkDBPγ 5’ and 3’UTRs using primers olFM245 and olFM0246 (adding SacII /SpeI sites) and primers olFM0247 and olFM248 (adding NotI/NcoI sites) respectively. A spacer sequence, to aid in subsequent diagnostic PCRs was generated by polymerase cycling assembly (PCA). Briefly, the spacer sequence was synthesised by using primers of 60 bp length with 20 bp homologous sequence to the adjacent primers on each side. Final concentrations of 0.6 μM for outer primers (ol488 and ol492) and 0.03 μM of inner primers (ol489, ol490, ol491 and ol503 were used for PCA with the same cycle conditions as described for PCR. The final product was inserted with SpeI and NcoI restriction sites between the HRs as described for pDonor_ *pkdbpα* cloning, to replace the deleted DBPγ genes. Primer sequences are listed in Table S4 and S6.

### Three-step nested PCR

Generation of each PCR repair template was carried out by a three-step nested PCR method to fuse together HRs with the insert DNA (eGFP expression cassette, eGFP with N-terminal linker or mCherry with C-terminal linker). In a first set of PCRs, the DNA insert (eGFP expression cassette or tag) and the HRs for integration into the region of interest were individually amplified in duplicate. The HRs contained at least 20 bp and 58°C Tm overhangs with homology to the insert DNA (HR1 with C-term overhang homologous to the N-term of insert DNA and HR2 with N-term overhang homologous to the C-term of the insert DNA). All duplicates were pooled and products were extracted from agarose gel (Qiagen) to remove primers and background amplicons. In a second nested PCR HR1 was fused to the donor amplicon in duplicate with double the amount of time allowed for the elongation step (10 s/kb) and again the product was gel extracted. In the final step the HR1-insert and HR2 were fused together resulting in the final product HR1-insert-HR2 (Fig 2A). PCR repair templates for HA tagging were generated in a two-step PCR method. First the HRs were individually amplified with addition of 27 bp HA sequence overhangs on the 3’end of HR1 and the 5’end of HR2. In the second nested PCR HR1 and HR2 were fused.

All primers are listed in Table S4 and all primer combinations for each contruct are listed in Table S5. Six to eight 50 μl reactions of the final construct PCRs were pooled (300 to 400 μl final volume and 20 μg DNA), ethanol precipitated and resuspended into sterile TE buffer for transfection. DNA concentrations were determined using Nano-Drop and band intensity measurement with BioRad Image lab software.

### DNA analysis

Genomic DNA from transfected parasite lines was extracted (QIAGEN) and analysed by PCR with GoTaq Master Mix (Promega) using the following conditions: 3 min at 96 °C, then 30 cycles of 25 s at 96 °C, 25 s at 52 °C, and 1 min/kb at 64 °C.

### Western blotting

To detect tagged proteins of interest, soluble cell extracts were prepared by lysing Nycodenz-enriched schizonts in 0.15 % saponin. Parasite pellets were washed several times with cold PBS and centrifugation at 13,000 rpm for 3 minutes at 4 °C to remove haemoglobin and red cell debris. Pellets were lysed in 5 pellet volumes of RIPA buffer (25 mM Tris, 150 mM NaCl, 1 % Triton X-100, 0.5 % Sodium deoxycholate, 0.1 % SDS, pH 7.5, 1x Roche Protease Inhibitors) supplemented with 50 units BaseMuncher (Expedeon) on ice for 20 minutes. This whole cell lysate was clarified by centrifugation at 13,000 rpm for 30 minutes at 4 °C. Soluble extracts were separated on Mini-Protean 4-20 % TGX gels (Bio-Rad) and transferred to nitrocellulose using the Trans-blot Turbo system (Bio-Rad). Equivalent uninfected red cell lysate or wild type *P. knowlesi* schizont lysates were analysed alongside lysates containing tagged proteins of interest. Membranes were blocked overnight, and tagged proteins were detected with mouse anti-GFP (Sigma, 1:5,000), rat anti-HA (Sigma 3F10 clone, 1:5,000), or rabbit anti-mCherry (ChromoTek, 1:5,000). Primary antibodies were detected using HRP- conjugated secondary antibodies (Bio-Rad, 1:5,000) and ECL (ThermoFisher Pierce). Chemiluminescence was captured using the Azure c600 system.

### Immunofluorescence assays and live cell imaging

Immunofluorescence assays were performed using blood smears fixed with 4 % paraformaldehyde for 30 min followed by washing in PBS and permeabilisation in 0.1 % Triton-X100 for 10 min. Slides of HA-tagged parasite lines were blocked overnight at 4°C in 3 % bovine serum albumin/PBS and then labelled with rabbit anti-HA high affinity (1:250) and Alexa Fluor 488-conjugated α-rabbit IgG (1:5000) (Thermo Fisher Scientific). The smears were mounted in ProLong Antifade mountant with DAPI (Thermo Fisher Scientific). For live cell imaging parasites were stained with Hoechst 33342 (New England Biolabs), transferred to poly-L-lysine-coated μ-slides VI (Ibidi, Martinsried, Germany). Both live and fixed preparations were viewed with a Nikon Ti E inverted microscope using a 100x oil immersion objective and imaged with an ORCA Flash 4.0 CMOS camera (Hamamatsu). Images were acquired and processed using the Nikon Elements Advanced Research software package.

### Invasion assays

Purified schizonts were set up in technical duplicate cultures with human RBCs, at a 2 % hematocrit and ~1 % parasitemia in 24 well plates. Parasitemia was measured with a flow cytometry (FACS)-based assay before and after incubation at 37 °C in a gassed chamber for 24 h. Samples were fixed with 2 % paraformaldehyde (Sigma) and 0.2 % glutaraldehyde (Sigma) in PBS for 1 h at 4C, washed, permeabilized with Triton X-100, and then washed again before 1 h RNase (MP Biomedicals) treatment, staining with SYBR Green I (Life Technologies), and FACS analysis. The samples were analyzed on a Becton Dickenson LSR-II. Data were acquired using FACSDiva 6.1.3 software and analyzed using FlowJo_V10. Three independent experiments were carried out in *Macaca fascicularis* blood and eight independent experiments were carried out in human blood. Statistical analyses was carried out with unpaired t-test comparing mean of PvDBP^OR^/Δβ and PvDBP^OR^/ΔβΔγ lines against the respective PkDBP^OR^ control parasite lines.

### Growth inhibition activity assays

Assays of growth inhibition activity (GIA), in the presence of anti-PvDBP_RII antibodies, were carried out using total IgG purified from rabbit sera using protein G columns (Pierce). Immunisation of rabbits against PvDBP_RII (SalI) has been described previously (55). Purified IgG was buffer-exchanged into RPMI 1640 medium, concentrated using ultra centrifugal devices (Millipore) and filter sterilized through a 0.22 μm filter (Millipore) prior to being aliquoted and frozen at −20 °C until use.

*P. knowlesi* parasites were synchronized by magnetic separation (MACS LS columns, Miltenyi Biotech). Synchronized trophozoites were adjusted to 1.5 % parasitemia, and 20 μL aliquots were pipetted into 96-well flat/half area tissue culture cluster plates (Appleton Woods). 20 μL purified IgG were added to triplicate test wells at eight final concentrations (10, 5, 2.5, 1.25, 0.625, 0.312, 0.15 and 0.075 mg/mL) and incubated for one cycle (26-30 h). Parasitemia was measured using the lactate dehydrogenase (pLDH) activity assay following standard protocols (56). An anti-DARC Fy6 VHH nanobody (45), a kind gift from Dr Olivier Bertrand (INSERM, France), was included in the test plate as a positive control in every assay (final concentration 1.5 or 3 µg/mL) and purified control IgG from the pre-immunisation sera of matched rabbits were used as the negative control. Anti-PkMSP119 rabbit sera, a kind gift from Ellen Knuepfer (Crick Institute, UK), was also tested in a similar manner. GIA of the purified IgG was expressed as percent inhibition calculated as follows: 100 − [(OD650 of infected erythrocytes with test IgG − OD650 of normal erythrocytes only) / (OD650 of infected erythrocytes without any IgG − OD650 of normal erythrocytes only) x 100 %].

### Whole genome sequencing

Genomic DNA was prepared for the PvDBP^OR^/ΔβΔy using the Blood and Tissue Kit (Qiagen). DNA libraries were prepared using the QIAseq FX DNA Library Kit (Qiagen) as per manufacturer’s instructions. A 20-minute fragmentation step was optimized for *Plasmodium* samples. Whole genome sequencing was performed using Illumina MiSeq technology with 150-base paired end fragment sizes. Raw sequence data for the A1-H.1 parental line was extracted from the European Nucleotide Archive as per (33, 57). The raw sequence data (accession number ERS3042513) was processed as previously described (58). In brief, the raw sequence data was aligned onto the A1-H.1 reference genome using the *bwa*-*mem* short read alignment algorithm (59), and coverage statistics were obtained using the *sambamba* software (60) to be plotted using R.

## Acknowledgements

This work is supported by an MRC Career Development Award (MR/M021157/1) jointly funded by the UK Medical Research Council and Department for International Development (R.W.M, F.M). M.N.H. is supported by a Bloomsbury Colleges research studentship. J.A.C. is supported by a BBSRC LIDO studentship. T.A.R. held a Wellcome Trust Research Training Fellowship [108734/Z/15/Z]; T.G.C. is funded by the Medical Research Council UK [ref. MR/M01360X/1, MR/N010469/1, MR/R025576/1, and MR/R020973/1] and BBSRC [ref. BB/R013063/1]. S.C. is funded by Medical Research Council UK grants [ref. MR/M01360X/1, MR/R025576/1, and MR/R020973/1]. S.J.D. is a Jenner Investigator, a Lister Institute Research Prize Fellow and a Wellcome Trust Senior Fellow [106917/Z/15/Z]. We are grateful to the Wellcome Trust for a Senior Investigator Award to D.A.B. (ref. 106240/Z/14/Z). R.C.H. is supported by a Marshall Scholarship granted by Her Majesty’s Government. C.J.S. is supported by Public Health England.

We thank Jose-Juan Lopez-Rubio for providing the pUF and pl6 plasmid, and also David Llewellyn, Jennifer Marshall and Doris Quinkert (University of Oxford) for assistance with rabbit immunisations, parasite culturing and the assays of GIA. We would also like to thank Michael Blackman (Francis Crick Institute) for providing Compound 2.

## Author Contributions

F.M. and M.N.H. designed and generated PCR constructs, performed parasite transfections, growth assays and microscopy. F.M., M.N.H. and A.P designed and generated plasmid constructs. T.A.R. carried out GIA assays and R.H. carried out Western blots. J.A.C and E.D.B carried out genome sequence analysis. J.H., and N.A. contributed new reagents, R.W.M, S.J.D., D.B, C.J.S, F.M., S.C, T.G.C and T.R. contributed to study design. F.M., R.H. and R.W.M wrote the paper. All the authors discussed the results and commented on the manuscript.

## Supplemental Figure legends

**Figure S1:**
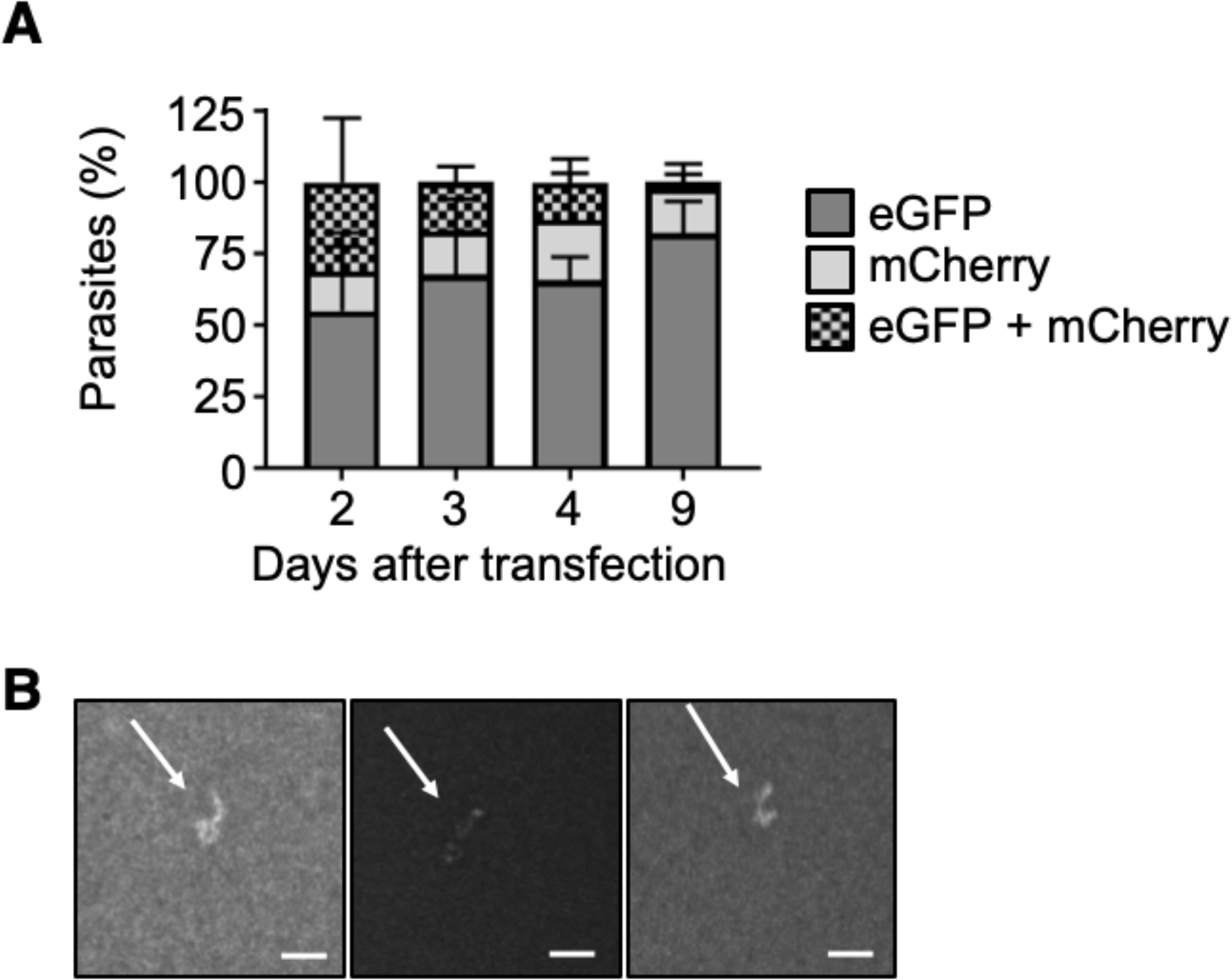
*P. knowlesi* dual plasmid uptake and plaque assay. (A) *P. knowlesi* parasites were co-transfected with 20 μg of plasmid containing an eGFP expression cassette (PkconGFP) and 20 μg of plasmid containing an mCherry expression cassette (PkconmCherry) and the proportion of parasites expressing each fluorescent protein monitored on consecutive days after transfection. Graph shows mean proportion of parasites expressing eGFP, mCherry or both across three independent experiments. Error bars denote ± 1 SD. (B) *P. knowlesi* parasites modified using CRISPR Cas9 were cloned by limiting dilution in 96 well plates. Infected wells were identified by scanning for parasite “plaques” 10 days after initiating cloning plates. Images show three representative images of parasite plaques visualised using 4X objective of an inverted microscope. Scale bars indicate 200 μm.

**Figure S2:**
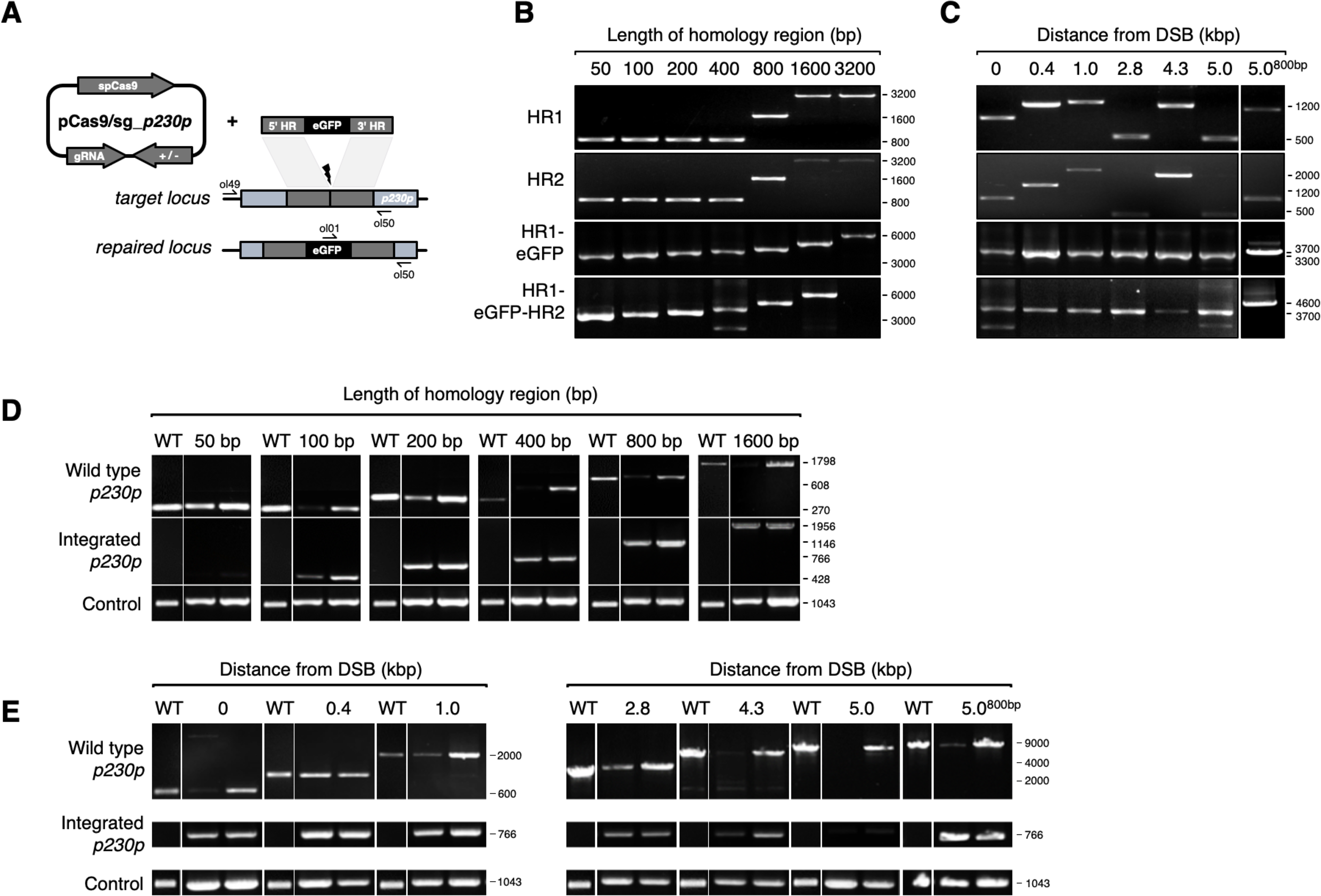
Schematic and genotypic analysis of fusion PCR repair template integration into *p230p* locus. (A) Schematic of CRISPR-Cas9 strategy. Integration of the eGFP expression cassette into the target *p230p* locus via homologous recombination using PCR repair template. Arrows indicating oligo positions for diagnostic PCRs. (B) Three-step nested PCR products to generate CRISPR-Cas9 repair templates. In the first step Homology region 1 (HR1) and HR2 are generated individually with eGFP cassette adaptors. Varying band sizes (e.g. for 50 to 400 bp HRs the same 800 bp product was generated) reflect different length of overhangs to allow a subsequent nested PCR step. In the second step HR1 and eGFP cassette are fused followed by the third and final step of fusing HR2 to the previous generated HR1-eGFP product. (C) Parasites transfected with pCas9/sg_*p230p* and PCR repair templates with HRs ranging from 50 to 1600 bp were analysed with diagnostic PCRs. PCRs of two transfections are shown. PCR reactions detecting the wild type locus, integration locus and a control PCR targeting an unrelated locus (ol75+ol76) using approximately 3 ng/µl genomic DNA for each reaction. (D) Parasites transfected with pCas9/sg_*p230p* and PCR repair templates with HRs of 400 bp or 800 bp of varying distances to the DSB were analysed with diagnostic PCRs. PCRs of two transfections are shown. PCR reactions detecting the wild type locus, integration locus and a control PCR targeting an unrelated locus (ol75+ol76) using approximately 3 ng/µl genomic DNA. Primers are listed in Table S4.

**Figure S3:**
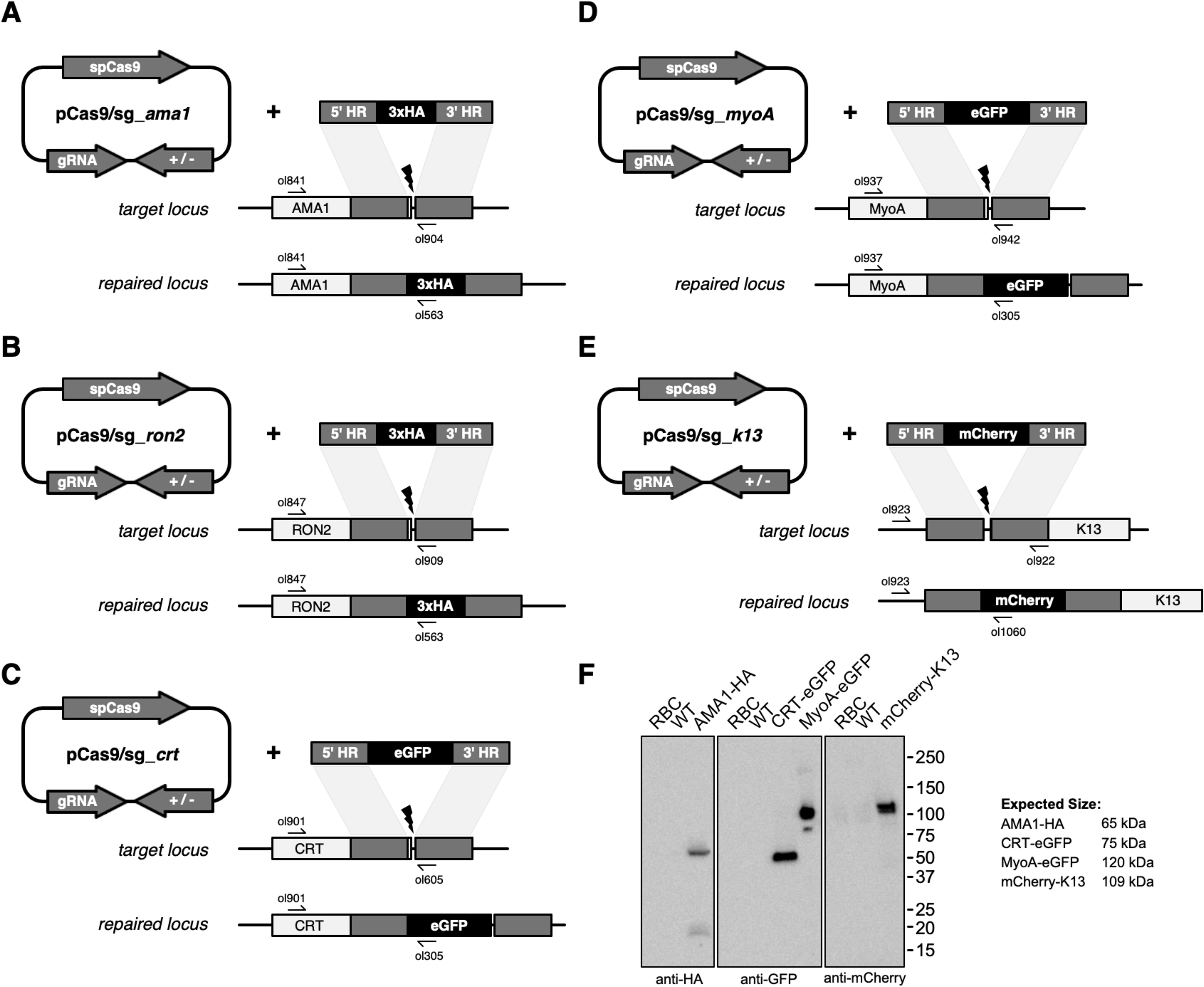
CRISPR-Cas9 tagging of *P. knowlesi* proteins. (A) Schematic of CRISPR-Cas9 strategy. C-terminal integration of the hemagglutinin (HA) tag into the target AMA1 locus via homologous recombination. Arrows indicating oligo positions for diagnostic PCRs. (B) C-terminal integration of HA tag into the target RON2 locus. (C) C-terminal integration of eGFP into the target Chloroquine Resistance Transporter (CRT) locus. (D) C-terminal integration of eGFP into the target Myosin A locus. (E) N-terminal integration of mCherry into the target Kelch13 (K13) locus. (F) Western blot showing expression of tagged proteins in *P. knowlesi* saponin-lysed schizonts separated by SDS-PAGE and immunoblotting with anti-HA, anti-GFP or anti-mCherry primary antibodies. Control samples are saponin-lysed red blood cells (RBC) and wild type parasite line. Expected sizes of bands are indicated. CRT runs faster than its expected band size as shown in *P. falciparum* previously.

**Figure S4:**
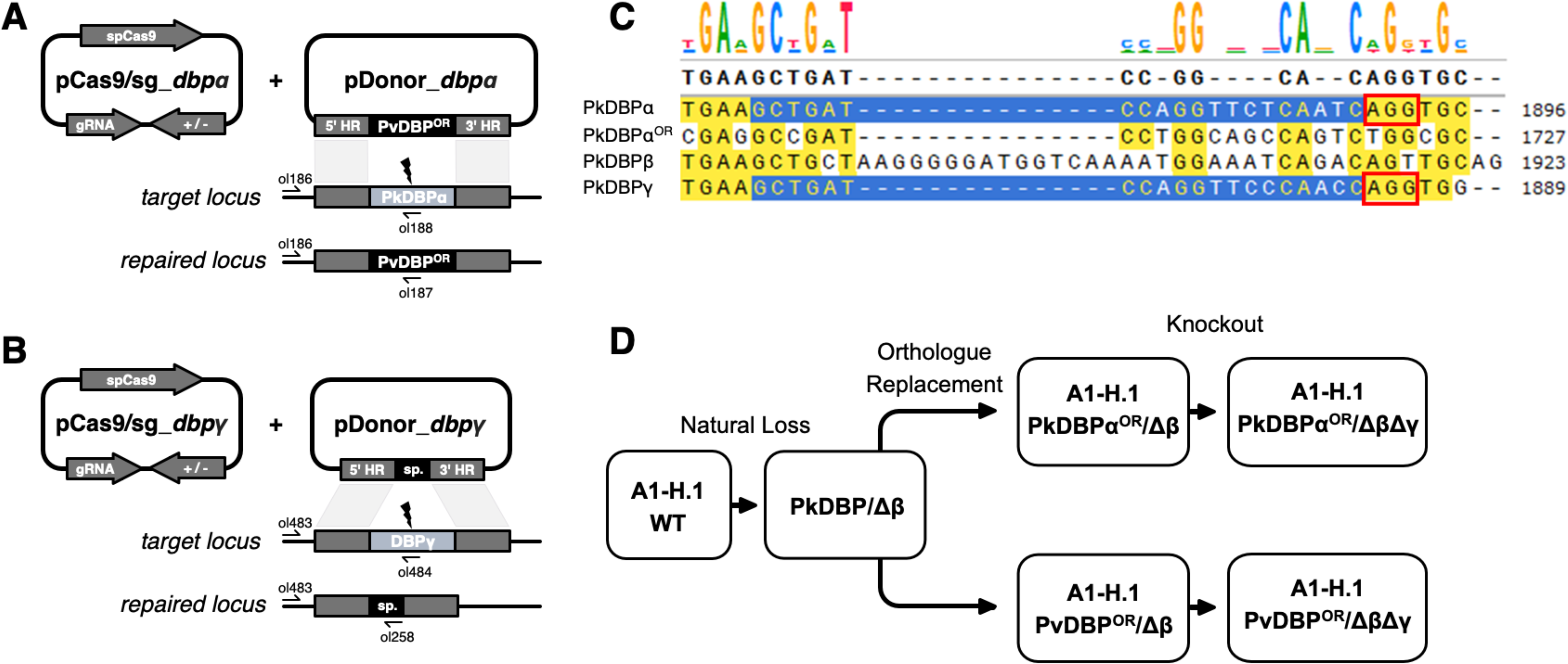
Transgenic *P. knowlesi* DBP orthologue replacement and knockout design and genotypic analysis. (A) Schematic of CRISPR-Cas9 strategy. Integration of the PvDBP into the target PkDBPα locus via homologous recombination. Arrows indicating oligo positions for diagnostic PCRs. (B) Integration of a spacer sequence into the target PkDBPγ locus. (C) Nucleotide sequence alignment of PkDBPα, PkDBPα^OR^, PkDBPβ and PkDBPγ guide sequences. The chosen guide sequences for transfections are highlighted in blue. PAM sites are highlighted in a red square (D) Flow chart indicating generated transgenic DBP parasite lines. Wild type parasites that naturally lost 44 kb at one end of chromosome 14 were first edited by orthologous replacement of PkDBPα with the recodonised gene (PkDBPα^OR^) or *P. vivax* DBP (PvDBP^OR^) and clonal lines established by limiting dilution cloning. Both clonal parasite lines were edited by knockout of PkDBPγ (PkDBPα^OR^/ΔβΔγ and PvDBP^OR^/ΔβΔγ), which were cloned before use in invasion assays and assays of GIA.

**Figure S5:**
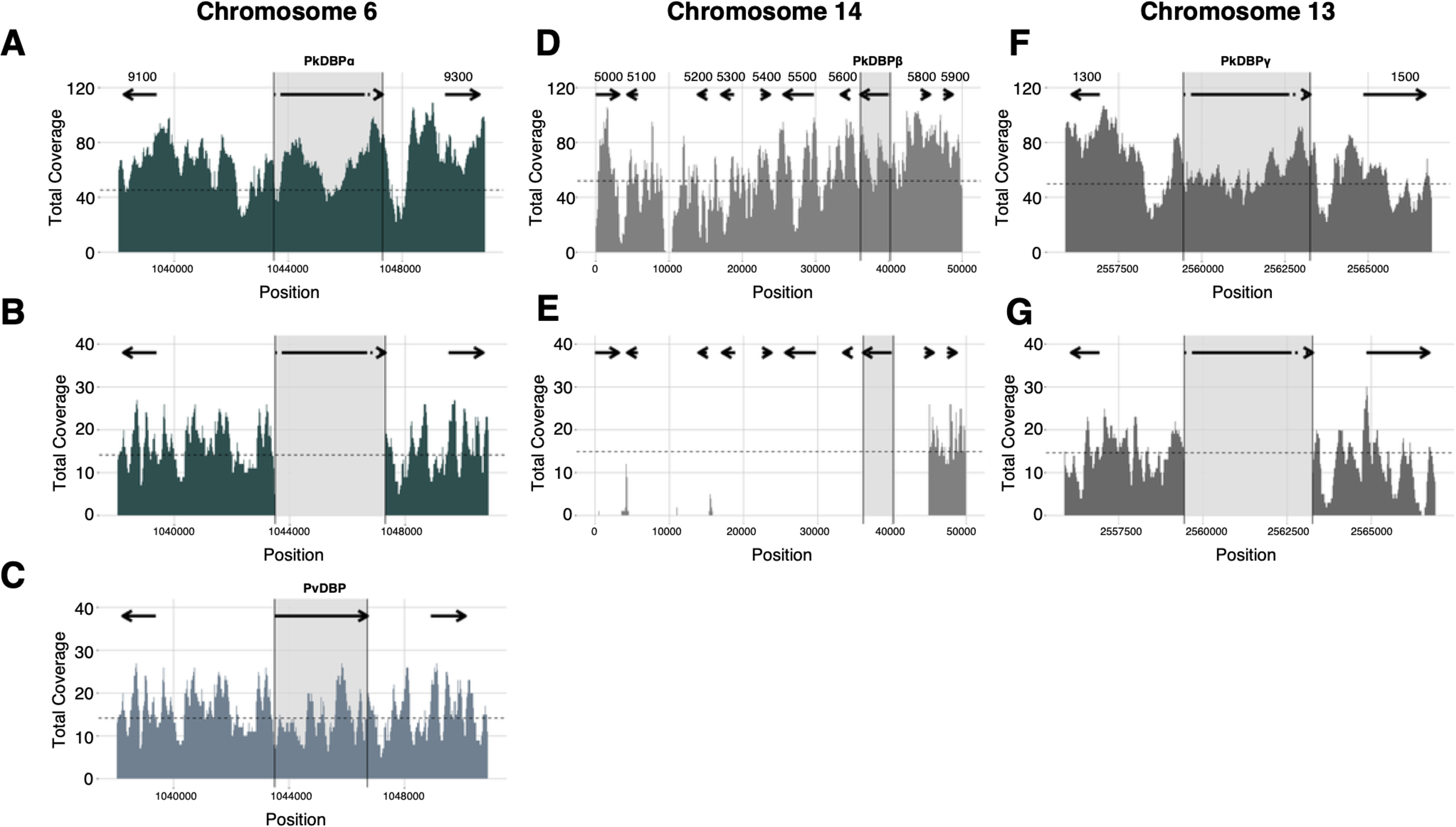
Sequencing of PvDBP^OR^/ΔβΔγ parasite line. Mapping of Illumina reads of PvDBP^OR^/ΔβΔγ against *P. knowlesi* strain A1-H.1 wild-type chromosomes (57). (A) PkDBPα locus on end of chromosome 6 of the A1-H.1 reference genome with flanking genes PKA1H_060029100 and PKA1H_060029300. (B) PvDBP^OR^/ΔβΔγ sequence mapped to A1-H.1 shows deletion of PkDBPα. (C) PvDBP^OR^/ΔβΔγ sequence mapped to a chimaeric A1-H.1 reference genome, generated *in silico* by replacing PkDBPα with PvDBP, confirms successful orthologue replacement (D) PkDBPβ locus of the A1-H.1 reference genome with flanking genes on start of chromosome 14 with flanking genes from PKA1H_140005000 to PKA1H_140005900, (E) PvDBP^OR^/ΔβΔγ sequence mapped to A1-H.1 reveals a chromosome truncation of 44,921 bp, including loss of PkDBPβ and 8 other genes. (F) PkDBPγ locus on end of chromosome 13 of the A1-H.1 reference genome with flanking genes PKA1H_130061300 and PKA1H_130061500. (G) PvDBP^OR^/ΔβΔγ sequence mapped to A1-H.1 shows deletion of DBPγ. Gene locations are indicated by arrows and last four digits of accession numbers are shown in the top panel above the arrows.

**Table S1:** Comparison of P. knowlesi and P. falciparum 3D7 genes.

| Gene | GC content for 5' and 3' HR regions |  | PAM site crossing stop codon<br>(off-target score) |  |
| --- | --- | --- | --- | --- |
|  | *P. knowlesi* | *P. falciparum* | *P. knowlesi* | *P. falciparum* |
| AMA1 | 43% / 36 % | 30 % / 13% | 2 sites | 1 site (0.032) |
| RON2 | 39 % / 36 % | 30 % / 13 % | 3 sites | no site |
| Myosin A | 41 % / 38 % | 29 % / 15 % | 1 site (not used) | no site |
| K13 | 34 % / 32 % | 8 % / 22 % | 3 sites | 2 sites, (0.057, 0.042) |
| CRT | 40 % / 36 % | 19 % / 10 % | 1 site | no site |

**Table S2:**
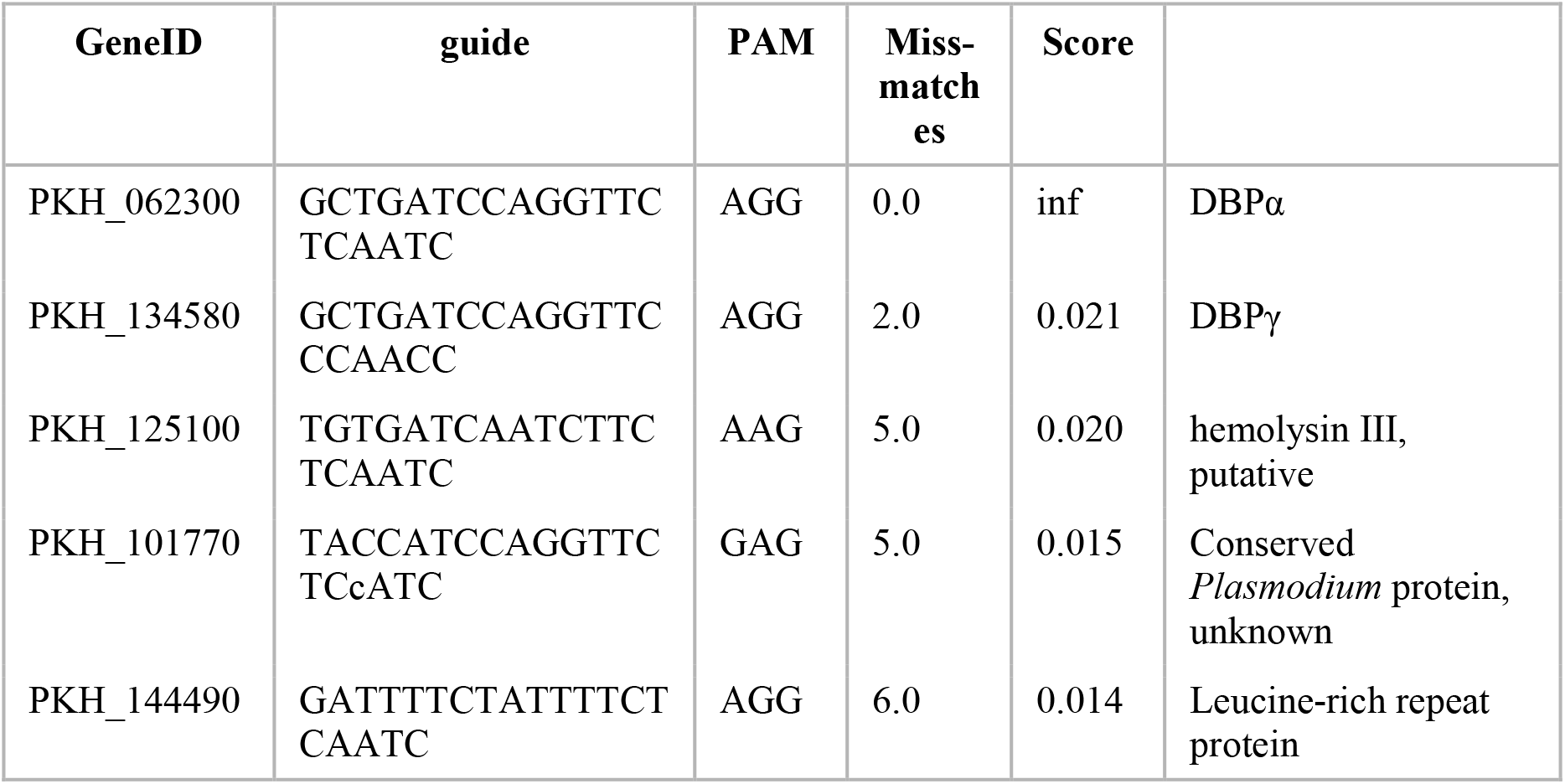
off-target scores of guide sequences for PkDBPα sgRNA.

**Table S3:**
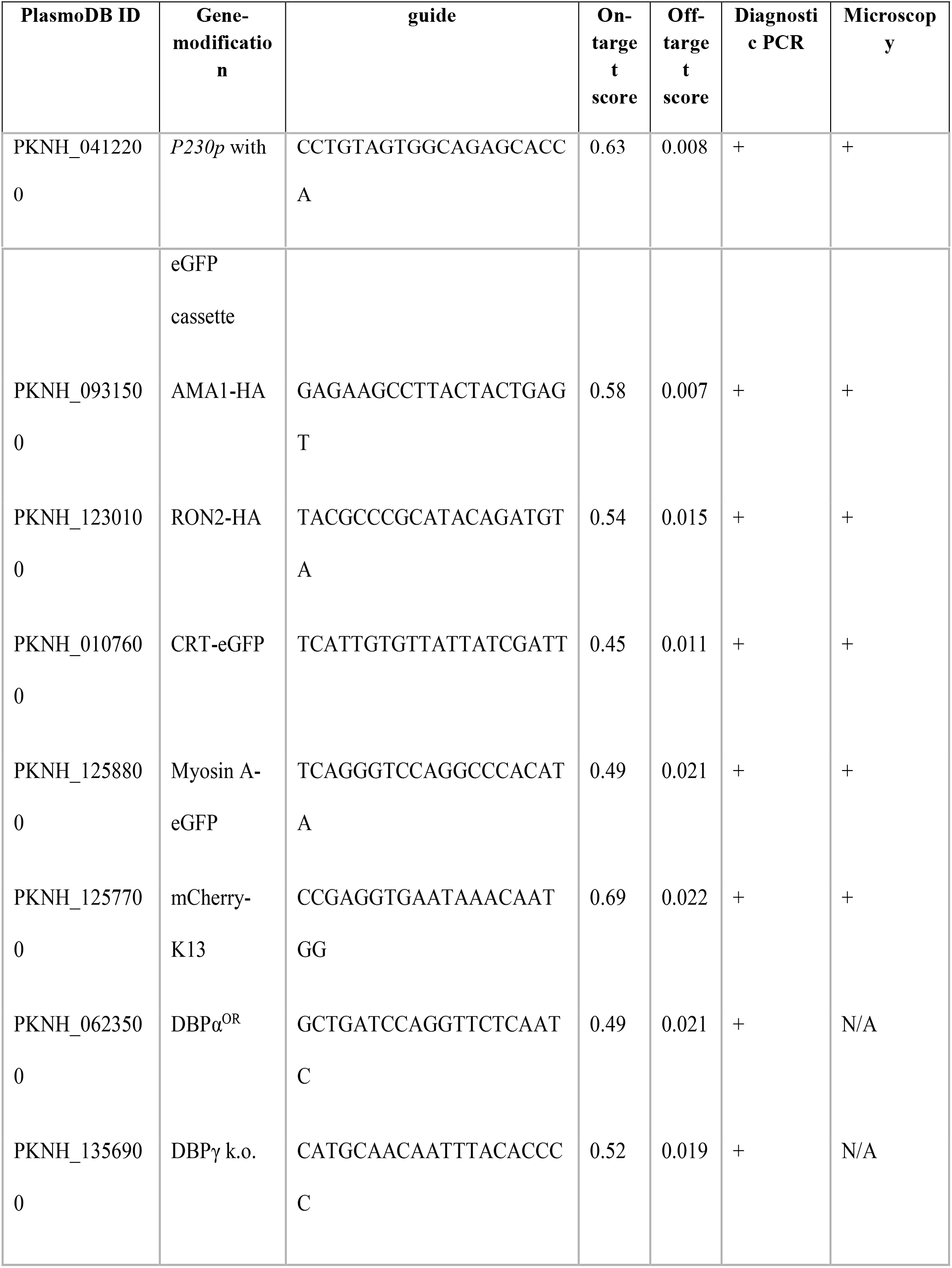
Guide sequences.

**Table S7:**
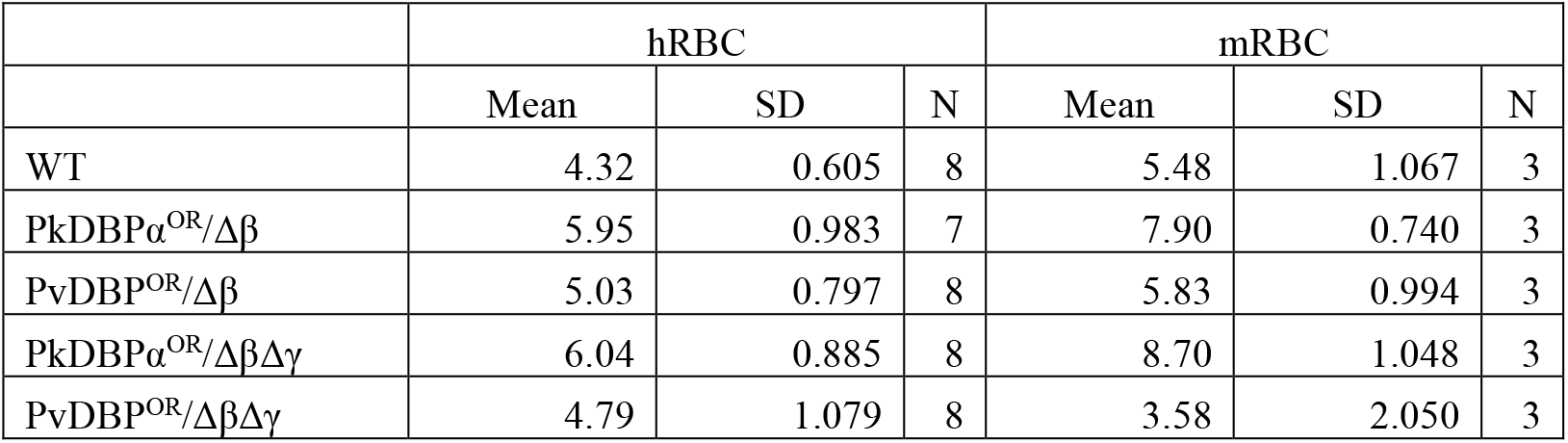
Fold replication of parasites lines in a FACS-based invasion assays over one growth cycle (24 h). Dataset of Fig 4B.

**Table S8:**
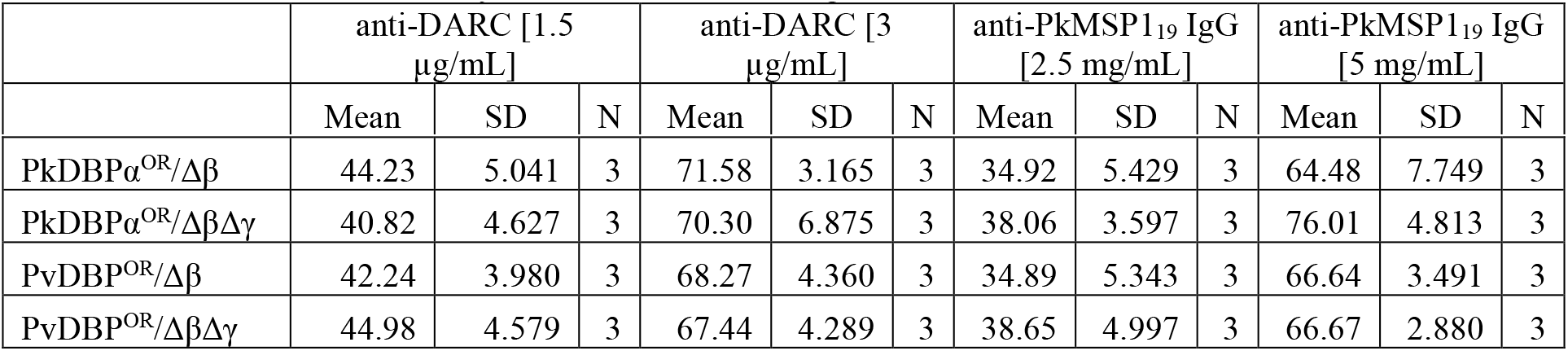
Growth inhibition activity (GIA, %). Dataset of Fig 4C.

| | anti-DARC [1.5 $\mu\text{g/mL}$] | | | anti-DARC [3 $\mu\text{g/mL}$] | | | anti-PkMSP1<sub>19</sub> IgG [2.5 mg/mL] | | | anti-PkMSP1<sub>19</sub> IgG [5 mg/mL] | | |
| --- | --- | --- | --- | --- | --- | --- | --- | --- | --- | --- | --- | --- |
|  | Mean | SD | N | Mean | SD | N | Mean | SD | N | Mean | SD | N |
| PkDBP$\alpha^{OR}/\Delta\beta$ | 44.23 | 5.041 | 3 | 71.58 | 3.165 | 3 | 34.92 | 5.429 | 3 | 64.48 | 7.749 | 3 |
| PkDBP$\alpha^{OR}/\Delta\beta\Delta\gamma$ | 40.82 | 4.627 | 3 | 70.30 | 6.875 | 3 | 38.06 | 3.597 | 3 | 76.01 | 4.813 | 3 |
| PvDBP$^{OR}/\Delta\beta$ | 42.24 | 3.980 | 3 | 68.27 | 4.360 | 3 | 34.89 | 5.343 | 3 | 66.64 | 3.491 | 3 |
| PvDBP$^{OR}/\Delta\beta\Delta\gamma$ | 44.98 | 4.579 | 3 | 67.44 | 4.289 | 3 | 38.65 | 4.997 | 3 | 66.67 | 2.880 | 3 |

**Table S9:**
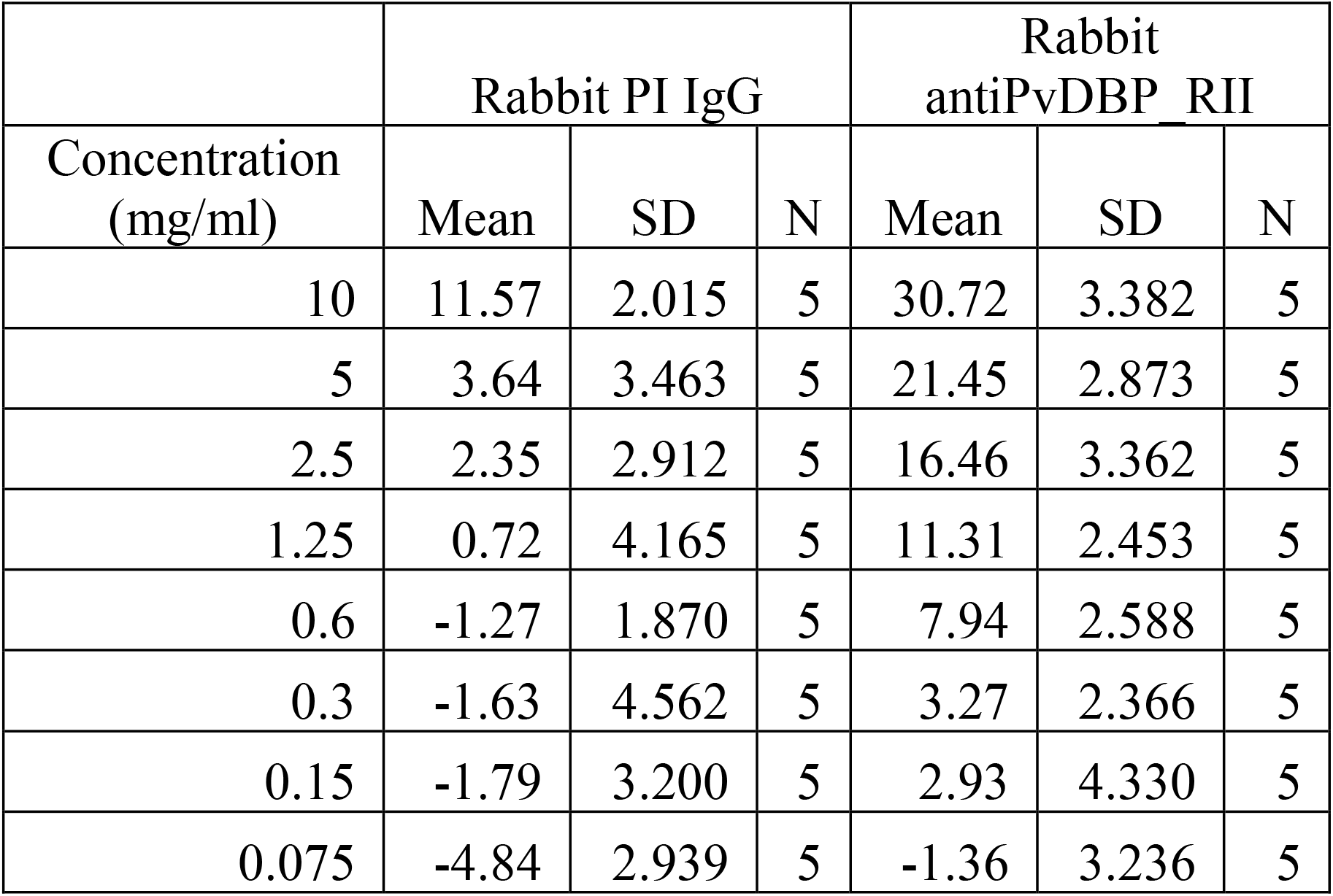
Growth inhibition activity (GIA, %) against parasite line PkDBPα^OR^/Δβ. Dataset of Fig 4D.

**Table S10:** Growth inhibition activity (GIA, %) against parasite line PvDBP^OR^/Δβ. Dataset of Fig 4E.

| Concentration<br>(mg/ml) | Rabbit PI IgG |  |  | Rabbit<br>antiPvDBP_RII |  |  |
| --- | --- | --- | --- | --- | --- | --- |
|  | Mean | SD | N | Mean | SD | N |
| 10 | 20.03 | 3.026 | 5 | 76.76 | 5.219 | 5 |
| 5 | 10.80 | 3.444 | 5 | 53.83 | 4.623 | 5 |
| 2.5 | 3.34 | 2.749 | 5 | 29.58 | 3.332 | 5 |
| 1.25 | 1.08 | 2.051 | 5 | 18.99 | 3.367 | 5 |
| 0.6 | 0.47 | 2.963 | 5 | 13.61 | 4.163 | 5 |
| 0.3 | -0.76 | 3.011 | 5 | 11.68 | 2.352 | 5 |
| 0.15 | -1.71 | 3.709 | 5 | 7.46 | 2.428 | 5 |
| 0.075 | -3.39 | 2.554 | 5 | -0.98 | 1.804 | 5 |

**Table S11:** Growth inhibition activity (GIA, %) against parasite line PkDBPα^OR^/ΔβΔγ. Dataset of Fig 4F.

| Concentration<br>(mg/ml) | Rabbit PI IgG |  |  | Rabbit<br>antiPvDBP_RII |  |  |
| --- | --- | --- | --- | --- | --- | --- |
|  | Mean | SD | N | Mean | SD | N |
| 10 | 28.70 | 2.828 | 6 | 14.18 | 3.589 | 6 |
| 5 | 20.31 | 2.385 | 6 | 4.34 | 3.060 | 6 |
| 2.5 | 15.45 | 2.723 | 6 | 3.17 | 2.337 | 6 |
| 1.25 | 10.96 | 2.270 | 6 | 0.32 | 1.661 | 6 |
| 0.6 | 7.39 | 3.215 | 6 | -0.03 | 2.501 | 6 |
| 0.3 | 5.90 | 3.156 | 6 | -2.28 | 2.374 | 6 |
| 0.15 | 3.05 | 3.498 | 6 | -1.21 | 1.559 | 6 |
| 0.075 | 0.90 | 2.428 | 6 | -0.77 | 2.185 | 6 |

**Table S12:** Growth inhibition activity (GIA, %) against parasite line PvDBP^OR^/ΔβΔγ. Dataset of Fig 4G.

| Concentration<br>(mg/ml) | Rabbit PI IgG |  |  | Rabbit<br>antiPvDBP RII |  |  |
| --- | --- | --- | --- | --- | --- | --- |
|  | Mean | SD | N | Mean | SD | N |
| 10 | 75.12 | 4.202 | 6 | 22.33 | 3.037 | 6 |
| 5 | 50.96 | 2.753 | 6 | 14.69 | 3.371 | 6 |
| 2.5 | 24.58 | 2.697 | 6 | 10.01 | 2.727 | 6 |
| 1.25 | 17.85 | 2.752 | 6 | 7.21 | 3.096 | 6 |
| 0.6 | 15.21 | 3.014 | 6 | 1.31 | 2.945 | 6 |
| 0.3 | 11.38 | 2.324 | 6 | 0.76 | 2.255 | 6 |
| 0.15 | 4.46 | 1.767 | 6 | 0.19 | 1.971 | 6 |
| 0.075 | 0.55 | 2.666 | 6 | -3.00 | 2.888 | 6 |

